# Non-canonical Oncostatin M Signaling Provides Protection during Respiratory Viral and Bacterial infections

**DOI:** 10.64898/2026.07.30.741615

**Authors:** Yewoo Lee, Lilly J. Patneaude, Kevyn R. Martins, Riley M.F. Pihl, Ernest L. Dimbo, Marta Pérez-Vázquez, Archana Jayaraman, Erin Crossey, Elise M.R. Armstrong, Bradley E. Hiller, Isabella Garza, Markus Bosmann, Matthew R. Jones, Joseph P. Mizgerd, Lee J. Quinton, Katrina E. Traber

## Abstract

Pneumonia remains a major global health burden, highlighting the need for host-directed therapies to complement antimicrobial treatment. Here, we identify Oncostatin M (OSM) as a critical regulator of pulmonary host responses during influenza and bacterial pneumonia. Loss of OSM shifted lung macrophages toward a pro-inflammatory phenotype during influenza infection and exacerbated lung injury during bacterial pneumonia, demonstrating an essential role for OSM in limiting immunopathology. Unexpectedly, OSM induced Signal Transducer and Activator of Transcription 3 (STAT3) activation in the absence of the canonical OSM receptor subunit OSMrβ, revealing previously unrecognized non-canonical OSM signaling in the mouse lungs. Consistent with this finding, loss of OSMrβ did not phenocopy the severe disease observed with loss of OSM. Together, these findings identify OSM as a key regulator of pulmonary immunity and reveal unexpected complexity in OSM signaling during pneumonia.

## Introduction

Pneumonia remains a leading cause of morbidity and mortality globally.^1, 2, 3^ Although microbe-directed therapies remain the main treatment for pneumonia, it does not fully address the dysregulated host response that contributes substantially to disease severity and poor clinical outcomes.^4^ Pneumonia disproportionately affects individuals with impaired or underdeveloped immune responses, including immunocompromised patients,^5^ young children whose immune systems are still maturing,^6^ and the elderly who experience a decline in immune function.^7^ Moreover, the rapid rise of antimicrobial resistance is further diminishing the effectiveness of microbe-directed therapies.^8, 9^ Together, these challenges highlight the critical importance of the host immune response in determining disease outcome and underscore the need to better understand the mechanisms that regulate host defense to facilitate the development of effective host-directed therapies.^10, 11^

Oncostatin M (OSM) is an IL-6 family cytokine known for its diverse roles during health and disease,^12^ making it a potentially attractive therapeutic target for various diseases. In the lungs, OSM has been implicated in the pathogenesis of several diseases, including pulmonary fibrosis,^13, 14^ asthma,^15, 16^ lung cancer^17, 18, 19^ and recently, influenza.^20^ However, despite being consistently upregulated across multiple forms of pneumonia,^20, 21, 22, 23^ its role in this context is not fully understood.

OSM signals through the shared IL-6 receptor, glycoprotein 130 (gp130) dimerized with cytokine-specific receptors to create a signaling receptor complex that activates various downstream pathways including JAK/STAT, MAPK/ERK and PI3K/AKT pathways.^12, 24, 25,26^ In the lungs, OSM is mainly expressed by leukocytes and OSM receptor (OSMrβ) is expressed by structural cells, suggesting that OSM signals in a paracrine fashion.^20, 22, 27^ Since the discovery and cloning of mouse OSMrβ in 1998, it was thought that murine OSM can only signal through murine OSMrβ heterodimerized with gp130.^28, 29^ Furthermore, the AB loop within the helical bundle of mouse OSM is a key structural determinant of receptor specificity, promoting activation of OSMrβ while limiting signaling through leukemia inhibitory factor receptor (LIFrβ).^30^ Nevertheless, these studies were performed in mouse cell lines, primarily in the murine embryonic fibroblast-like NIH 3T3 cells and the murine alveolar macrophage-like MH-S cells, to assess mouse OSM-receptor interactions, leaving it unclear whether the findings are generalizable to other cell types. Using *in vivo* receptor knockdown approaches, studies using primary cells suggest that mouse OSM activates LIFrβ in certain contexts, such as mouse osteoblasts to regulate bone turnover, suggesting that non-canonical mouse OSM signaling promotes important biological function.^31, 32^ Whether non-canonical mouse OSM signaling occurs in the lungs is currently unknown.

In this study, we investigated the role of OSM in viral (influenza) and bacterial (*Escherichia coli* (*E. coli*)) pneumonia using mice deficient in the cytokine (OSM) or the receptor (OSMrβ). We discovered that non-canonical OSM signaling, independent of OSMrβ, is sufficient to protect against lung injury and excessive inflammation in influenza and bacterial pneumonia.

## Results

### OSM receptor independent OSM signaling is sufficient for protection against influenza

To investigate the role of OSM during influenza, we generated whole body OSM knockout mice (OSM^-/-^) and verified deletion of OSM in the lungs (Figure S1A). We confirmed that OSM is upregulated during influenza by intratracheally instilling a sublethal dose of X31 influenza into the left lobe of lungs of wildtype (WT) mice and measuring OSM RNA and protein (Figure S1B). We infected OSM^-/-^ and WT mice with influenza and observed that OSM^-/-^ mice exhibit greater morbidity compared to WT with increased weight loss starting at 5 days post-infection (Figure 1A). We found similar morbidity results with a lethal dose of influenza strain PR8 (Figure 1B). To investigate OSM-mediated immune responses without excessive mortality,^33^ we continued further studies using the less virulent X31 influenza strain.

**Figure 1:**
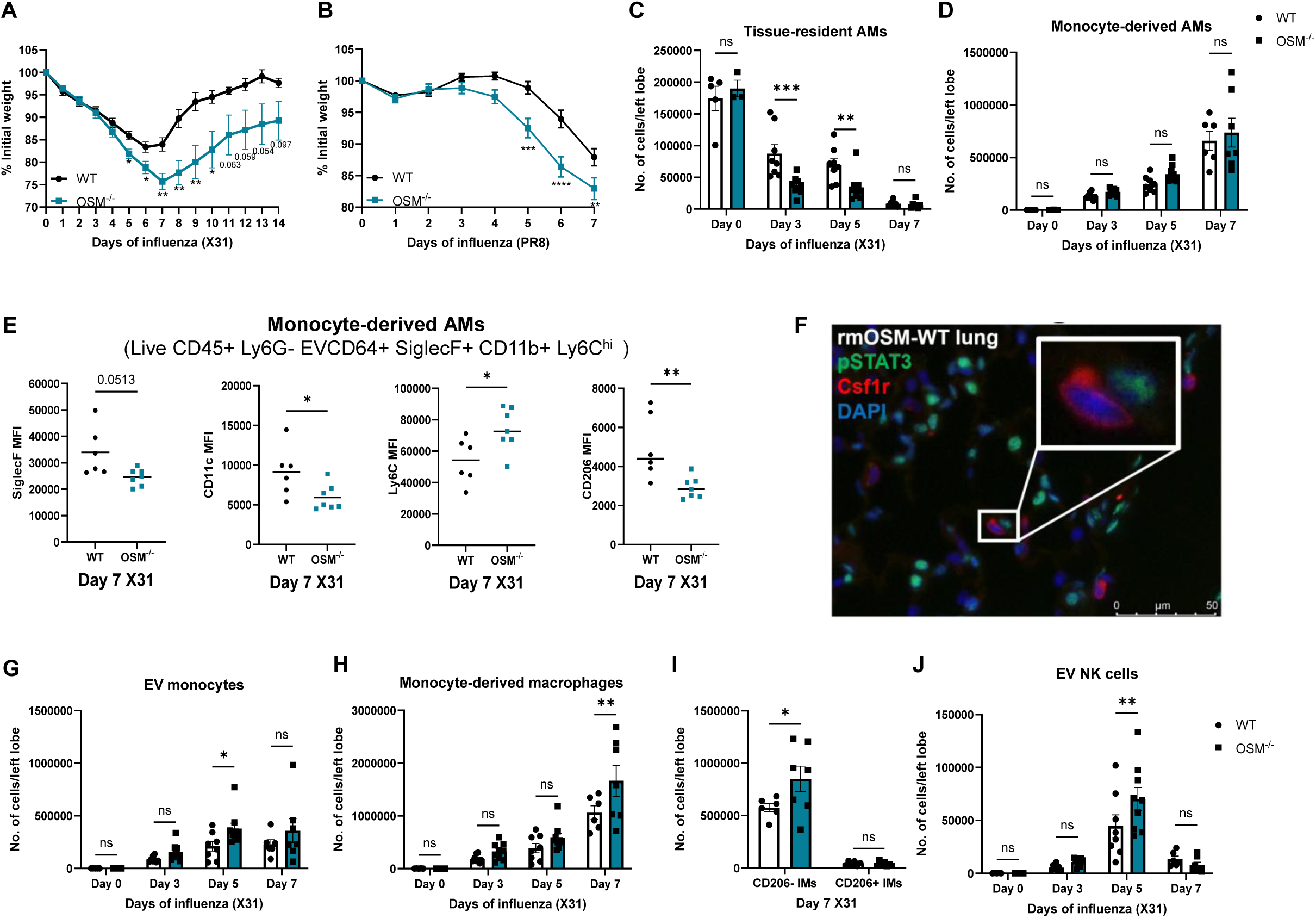
Loss of OSM increases morbidity and recruitment of inflammatory leukocytes during influenza infection. Weights of WT and OSM^-/-^ mice that were intratracheally infected with (A) 1000 plaque-forming units (PFU) of X31 (influenza A/HKx31, H3N2) (n=6 for WT and n=8 for OSM^-/-^) and (B) 200 PFU of PR8 (influenza A/Puerto Rico/8/34, H1N1) (n=20 for WT and n=19 for OSM^-/-^). Two-way ANOVA for (A) and (B) * P ≤ 0.05, ** P ≤ 0.01, *** P ≤ 0.001 and **** P < 0.0001. At least two independent experiments. Leukocytes in the left lung of WT and OSM^-/-^ mice harvested at baseline (day 0), day 3, day 5 and day 7 post X31 infection were quantified by spectral flow cytometry. Mice were intravascularly instilled with CD45.2 flow antibody to distinguish extravascular leukocytes from intravascular leukocytes. At least two independent experiments. (C) Absolute cell counts of EV (IV CD45.2-) tissue-resident alveolar macrophages, (D) absolute cell counts of EV (IV CD45.2-) monocyte-derived AMs (E) Median Fluorescence Intensity (MFI) of monocyte-derived AMs (F) IF of rmOSM treated WT lungs stained with pSTAT3 in green, Csf1r (myeloid marker) in red and DAPI nuclear stain in blue (G) absolute cell counts of EV (IV CD45.2-) monocytes, (H) absolute cell counts of EV (IV CD45.2-) monocyte-derived macrophages (I) absolute cell counts of EV (IV CD45.2-) CD206- and CD206+ inflammatory and interstitial macrophages (J) absolute cell counts of EV (IV CD45.2-) natural killer cells. Two-way ANOVA for (C-D, G-J) * P ≤ 0.05, ** P ≤ 0.01, *** P ≤ 0.001. Mann-Whitney for (E) * P ≤ 0.05, ** P ≤ 0.01. Each data point represents an individual mouse.

Given the increased morbidity observed with loss of OSM during influenza, we next sought to investigate the underlying mechanisms driving this phenotype. Because leukocyte responses play a central role in influenza pathogenesis,^34, 35^ we performed immunophenotyping of intravascular and extravascular leukocytes in the lungs at various timepoints of influenza to characterize changes in immune cell populations (gating strategy in Figure S2). As previously shown,^36, 37, 38, 39^ we observed decrease in tissue-resident alveolar macrophages (TRAMs, distinguished from monocyte-derived AMs (moAMs) as extravascular CD45^+^CD64^+^SiglecF^+^Ly6G^-^ cells lacking CD11b and Ly6C expression (Figure S2)) with influenza infection (Figure 1C). Interestingly, OSM^-/-^ lungs had greater reduction of TRAMs at 3 days and 5 days post-influenza infection compared to WT (Figure 1C). During influenza infection, Ly6C^+^ monocytes are recruited into the lungs and differentiate into AMs to replenish the reduced AM niche.^38, 40^ As the monocytes differentiate into AMs, they increase the expression of AM markers SiglecF and CD11c and decrease expression of monocyte marker Ly6C.^38^ Although we observed no difference in the number of moAMs with loss of OSM (Figure 1D), we saw decreased expression of AM markers SiglecF and CD11c and increased expression of monocyte marker Ly6C at day 7 post-influenza on moAMs in OSM^-/-^ lungs compared to WT (Figure 1E), suggesting the differentiation of monocyte to AM is slower in OSM^-/-^ lungs. We also observed a decrease in expression of CD206, a marker often associated with a more pro-resolving phenotype,^41, 42^ on moAMs in OSM^-/-^ lungs (Figure 1E). It has been shown that myeloid cells do not express OSMrβ.^20, 43, 44^ To determine whether OSM directly activates myeloid cells, we stimulated WT lungs with recombinant mouse OSM (rmOSM) and assessed STAT3 activation, a well-established downstream mediator of OSM signaling.^45^ We demonstrated that STAT3 is not activated in myeloid cells (Csf1r^+^) in WT lungs stimulated with rmOSM (Figure 1F). Hence, the macrophage changes observed following the loss of OSM are unlikely to result from a direct effect of OSM on macrophages. This finding is consistent with previous studies demonstrating that OSM does not directly activate leukocytes.^20, 21^

We noted an increase in extravascular monocytes 5 days post-infection in OSM^-/-^ lungs (Figure 1G) which may result in the increase in monocyte-derived macrophages (moMacs) at 7 days post-infection in OSM^-/-^ lungs compared to WT control (Figure 1H). MoMacs, which were further defined by high Ly6C expression (Figure S3), include inflammatory and interstitial macrophages (IMs) which comprise of at least two major subsets, CD206^+^ IMs which are located around the airways in the bronchial interstitium, and CD206^-^ IMs which are in the alveolar interstitium.^46, 47^ We observe that on day 7 post-influenza, CD206^-^ IMs were more abundant in the lung-derived cell suspensions than CD206^+^ IMs (Figure 1I). This distribution of IM subsets at this inflammatory timepoint during influenza infection is consistent with previously proposed functional heterogeneity, where CD206^-^IMs are often enriched for inflammatory and antigen-presentation programs, whereas CD206^+^ IMs are more commonly associated with immunoregulatory and tissue-homeostatic functions.^46, 48^ Interestingly, we observed an increase in the number of CD206^-^ IMs in OSM^-/-^ lungs compared to WT lungs but no difference in the number of CD206^+^ IMs (Figure 1I). We also observed an increase in NK cells at 5 days post-infection in OSM^-/-^ lungs (Figure 1J). Nevertheless, we did not observe any differences in neutrophils, eosinophils and Ly6G^+^ macrophages (Figure S4A-4C). There were also no differences in different types of dendritic cells (CD103^+^CD11b^-^, CD103^+^CD11b^+^ and CD103^-^CD11b^+^ conventional DCs and plasmacytoid DCs) between OSM^-/-^ lungs and WT control (Figure S4D-4G). Together, these findings indicate that loss of OSM during influenza most prominently impacts the macrophage compartment with a shift towards a proinflammatory phenotype with enhanced recruitment of monocytes and moMacs and reduced number of TRAMs.

In addition to leukocytes, accumulating evidence suggests that structural and stromal cells actively participate in protecting the host from invading pathogens during pneumonia.^11, 49 50^ We previously demonstrated that OSM signaling to lung epithelial cells regulates neutrophil recruitment through CXCL5 induction.^21^ To further investigate OSM signaling on lung epithelial cells during influenza we generated mice with targeted lung-epithelial OSMrβ deletion (EpiOSMrβ^-/-^). Epithelial-specific deletion of OSMrβ was validated by FACS isolation of epithelial cells and leukocytes, followed by immunoblot analysis demonstrating reduced OSMrβ expression in epithelial cells from Cre⁺ mice (Figure S5A). We infected EpiOSMrβ^-/-^ and WT mice with both sublethal (50PFU/left lobe) and lethal (400PFU/left lobe) doses of PR8. We observed no differences in mortality and morbidity (Figure S5B-5E), number of BALF leukocytes (Figure S5F-5G) and lung injury as measured by total BALF protein (Figure S5H). Taken together, this suggests that OSM-OSMrβ signaling on lung epithelial cells is not required for OSM mediated protection during influenza.

Because epithelial-specific OSM signaling was dispensable for OSM-mediated protection during influenza, we next investigated the contribution of global OSM-OSMrβ signaling to host defense during influenza. To study this, we generated whole body OSMrβ knockout mice (OSMrβ^-/-^) and confirmed deletion of OSMrβ protein from the whole lung (Figure S6A). We then infected OSM^-/-^, OSMrβ^-/-^ and WT control with sublethal dose of X31 influenza. As expected, OSM^-/-^ mice had greater morbidity and mortality compared to WT control. However, we observed no differences in morbidity and mortality between OSMrβ^-/-^ compared to WT control indicating that OSMrβ is not required for OSM mediated protection during influenza (Figure 2A-2B). We observed a similar pattern of weight loss with low dose (50PFU) of X31 influenza infection (Figure S6B).

**Figure 2:**
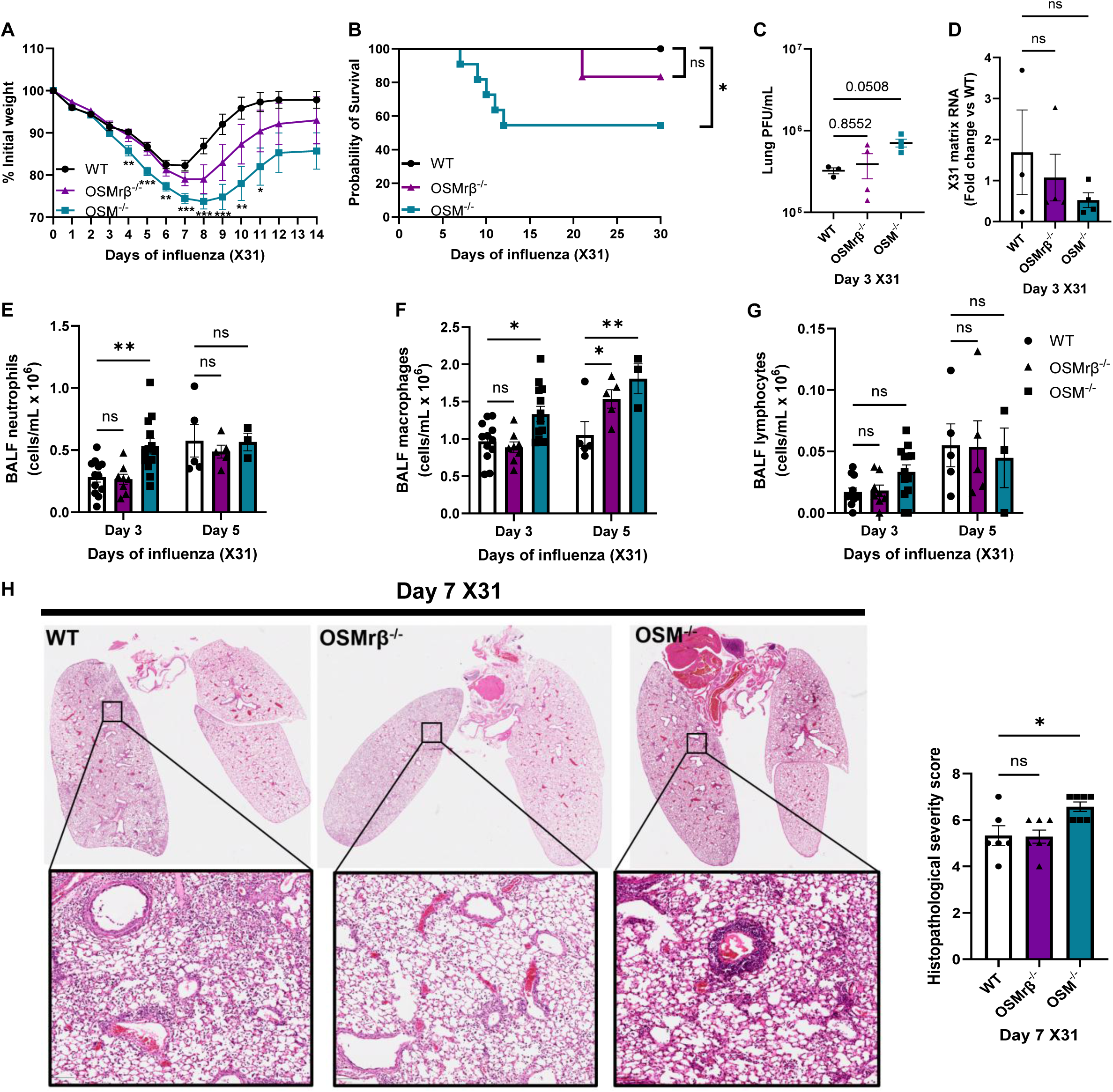
OSMrβ-independent OSM signaling is sufficient to mediate protection during influenza infection. (A) Weights of WT, OSMrβ^-/-^ and OSM^-/-^ mice that were intratracheally infected with 1000 PFU of X31 influenza (n=15 for WT, n=14 for OSMrβ^-/-^ and n=19 for OSM^-/-^). Mixed-effects analysis * P ≤ 0.05, ** P ≤ 0.01 and *** P ≤ 0.001. (B) Survival curve of WT, OSMrβ^-/-^ and OSM^-/-^ mice that were intratracheally infected with 1000 PFU of X31 influenza (n=8 for WT, n=6 for OSMrβ^-/-^and n=11 for OSM^-/-^). Log-rank (Mantel-Cox) test * P ≤ 0.05. Left lungs were collected at 3 days post-infection, and viral burden was determined either by (C) plaque assay to quantify plaque-forming units (PFU) or (D) by RT-qPCR measurement of X31 matrix transcript levels. One-way ANOVA. Number of (E) neutrophils, (F) macrophages and (G) lymphocytes in BALF of WT, OSMrβ^-/-^ and OSM^-/-^ lungs at 3 days and 5 days post X31 influenza. Two-way ANOVA for (E-G) * P ≤ 0.05, ** P ≤ 0.01. (H) WT, OSMrβ^-/-^ and OSM^-/-^ mice were intratracheally infected with 1000 PFU of X31 influenza virus. Lungs were harvested on day 7 post-infection, and histopathological severity was evaluated. Representative histological sections are shown. Kruskal-Wallis * P ≤ 0.05. All experiments were performed in at least two independent experiments. Each data point represents an individual mouse.

There was a mild increase in X31 plaque-forming units (PFU) in OSM^-/-^ lungs compared to WT control at 3 days post-influenza but no differences in viral load between OSMrβ^-/-^and WT control (Figure 2C). However, differences in X31 matrix RNA levels in the lungs were not present across the three groups (Figure 2D), suggesting that enhanced morbidity observed in OSM^-/-^ mice is not due to differences in viral burden. Differential cell counts by manual microscopic evaluation and cellular morphology were performed on cytocentrifuged BALF 3 and 5 days post-infection. OSM^-/-^ mice had an increase in BALF neutrophils on day 3 and macrophages on day 3 and 5 post-infection, most likely due to increase in monocytes and moMacs in OSM^-/-^ as observed in figure 1 (Figure 2E-2F). We also detected a slight increase in BALF macrophages in OSMrβ^-/-^ lungs compared to WT control at day 5 post-infection (Figure 2F). There was no difference in lymphocyte number between genotypes at both time points (Figure 2G). Consistent with the increased morbidity observed in OSM^-/-^ mice, histological analysis on day 7 post-influenza infection demonstrated significantly greater lung pathology in OSM^-/-^ mice compared with WT controls (Figure 2H). In contrast, OSMrβ^-/-^ mice displayed histopathological severity scores similar to those of WT controls (Figure 2H). In summary, our data suggest that OSMrβ-dependent signaling is dispensable for the protective effects of OSM during influenza infection, pointing to the existence of alternative signaling pathways that mediate OSM-driven host protection.

### OSM receptor independent OSM signaling is sufficient for protection against E. coli

To investigate the role of OSM during bacterial pneumonia, we intratracheally instilled *E. coli* into left lobe of lungs of OSM^-/-^, OSMrβ^-/-^, and WT mice. We have previously shown that neutralization of OSM leads to reduced neutrophil recruitment.^21^ We, however, observed no differences in BALF neutrophils and macrophages between OSM^-/-^ and WT mice at 6 hours, 24 hours and 48 hours of pneumonia (Figure 3A), nor did we observe any differences in BALF neutrophils and macrophages between OSMrβ^-/-^ and WT mice at 24 and 48 hours post-infection (Figure 3B). Since we cannot accurately distinguish between TRAMs and recruited monocytes using this method, we immunophenotyped *E. coli* infected OSM^-/-^ and WT lungs at 24 hours post-infection (gating strategy in Figure S7). Although there were no differences in the number of extravascular neutrophils and monocytes, we observed reduced TRAMs in OSM^-/-^ lungs (Figure 3C) similar to what we observed during influenza infection (Figure 1). We did not observe any differences in the number of TRAMs, extravascular neutrophils and monocytes in OSMrβ^-/-^ lungs compared to WT control (Figure 3D).

**Figure 3:**
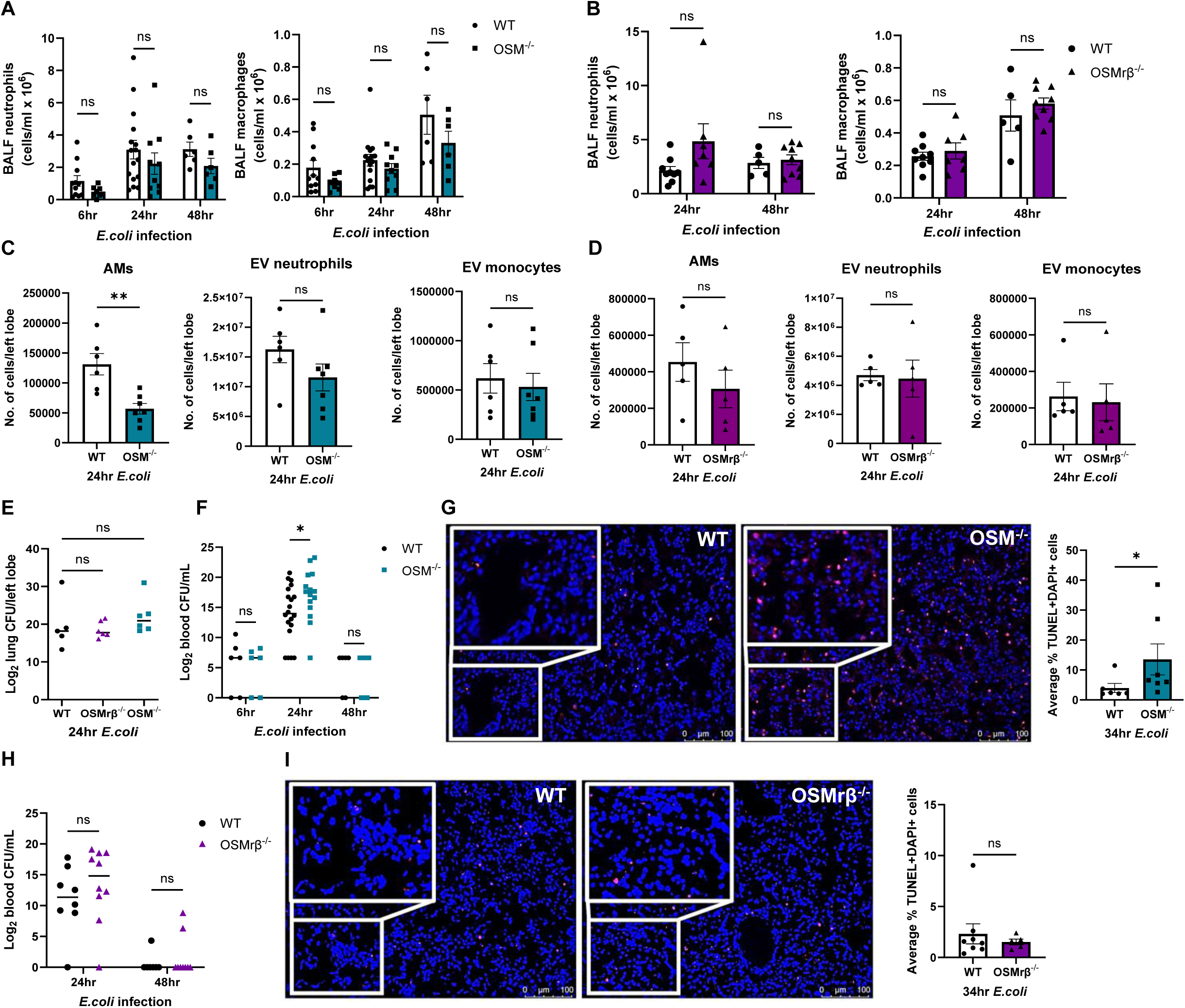
Loss of OSM but not OSMrβ leads to greater injury during *E.coli* pneumonia. (A) Number of neutrophils and macrophages in BALF of WT and OSM^-/-^ lungs at 6hr, 24hr and 48hr post *E.coli* pneumonia. (B) Number of neutrophils and macrophages in BALF of WT and OSMrβ^-/-^ lungs at 24hr and 48hr post *E.coli* pneumonia. Two-way ANOVA. Leukocytes in the left lung of mice at 24hr post *E.coli* pneumonia were quantified by spectral flow cytometry. Absolute cell counts of EV (IV CD45.2-) tissue-resident alveolar macrophages, neutrophils and monocytes in (C) WT and OSM^-/-^ lungs and (D) WT and OSMrβ^-/-^ lungs. Unpaired T-test ** P ≤ 0.01. (E) WT, OSMrβ^-/-^ and OSM^-/-^ mice were intratracheally infected with *E.coli.* Left lung was harvested at 24hr post-infection and bacterial burden was determined by quantifying colony-forming units (CFU). One-way ANOVA. (F) WT and OSM^-/-^ mice were intratracheally infected with *E.coli.* Blood was collected at 6hr, 24hr and 48hr post-infection and bacteremia was determined by quantifying CFU. Two-way ANOVA * P ≤ 0.05. (G) WT and OSM^-/-^ mice were intratracheally infected with *E.coli* and left lung was fixed at 34hr post-infection. Paraffin-embedded lung sections were stained for TUNEL (red) and DAPI (blue), and apoptotic cells were enumerated by microscopic analysis. Mann-Whitney * P ≤ 0.05. (H) WT and OSMrβ^-/-^ mice were intratracheally infected with *E.coli.* Blood was collected at 24hr and 48hr post-infection and bacteremia was determined by quantifying CFU. Two-way ANOVA. (I) WT and OSMrβ^-/-^ mice were intratracheally infected with *E.coli* and left lung was fixed at 34hr post-infection. Paraffin-embedded lung sections were stained for TUNEL (red) and DAPI (blue), and apoptotic cells were enumerated by microscopic analysis. Mann-Whitney. All experiments were performed in at least two independent experiments. Each data point represents an individual mouse.

Since loss of AMs is often associated with increased pathogen burden,^39, 51^ we then measured lung bacterial burden. We did not see any differences in lung CFU in OSM^-/-^lungs, OSMrβ^-/-^ lungs and WT controls at 24 hours post-infection (Figure 3E). However, we found a significant increase in bacteremia in OSM^-/-^ mice at 24 hours post-infection (Figure 3F). Since increased bacteremia is often associated with greater lung injury,^52, 53^ we measured cellular apoptosis in OSM^-/-^ and WT lungs by TUNEL staining at peak moribund timepoint of 34 hours. There was an increase in TUNEL-positive cells in OSM^-/-^ lungs, suggesting that the loss of OSM leads to greater injury (Figure 3G). However, we did not observe any differences in bacteremia and apoptosis in OSMrβ^-/-^ mice (Figure 3H-3I).

To see if there are any changes in downstream signaling pathways with the loss of OSM, we measured activation of transcription factors known to be regulated by OSM.^45^ We saw a decrease in STAT3 activation, trend towards decrease in STAT1 activation and increase in ERK activation in OSM^-/-^ lungs at 6 hours post-*E. coli* infection (Figure S8A-8C). We did not observe any changes in STAT3 activation in OSMrβ^-/-^ mice (Figure S8D). Taken together, these data suggest that OSM is required for protection against lung injury during *E. coli* pneumonia and that this protective effect can occur independently of canonical OSM-OSMrβ signaling.

### OSM signaling in the lungs occurs both in the presence and absence of OSMrβ

Our data so far suggest an unanticipated finding that canonical OSM-OSMrβ signaling is not required for OSM mediated protection during pneumonia. To determine if the lungs respond to OSM in the absence of OSMrβ, we intratracheally instilled rmOSM or vehicle control (PBS) for 1 hour into WT and OSMrβ^-/-^ lungs and measured downstream transcription factor activation by immunofluorescence staining. OSM induced STAT3 activation in both WT and OSMrβ^-/-^ lungs, while PBS treatment did not (Figure 4A). Similar observations were seen with immunoblotting analysis as well (Figure S9). These results suggest that OSM can activate the lungs even in the absence of OSMrβ.

**Figure 4:**
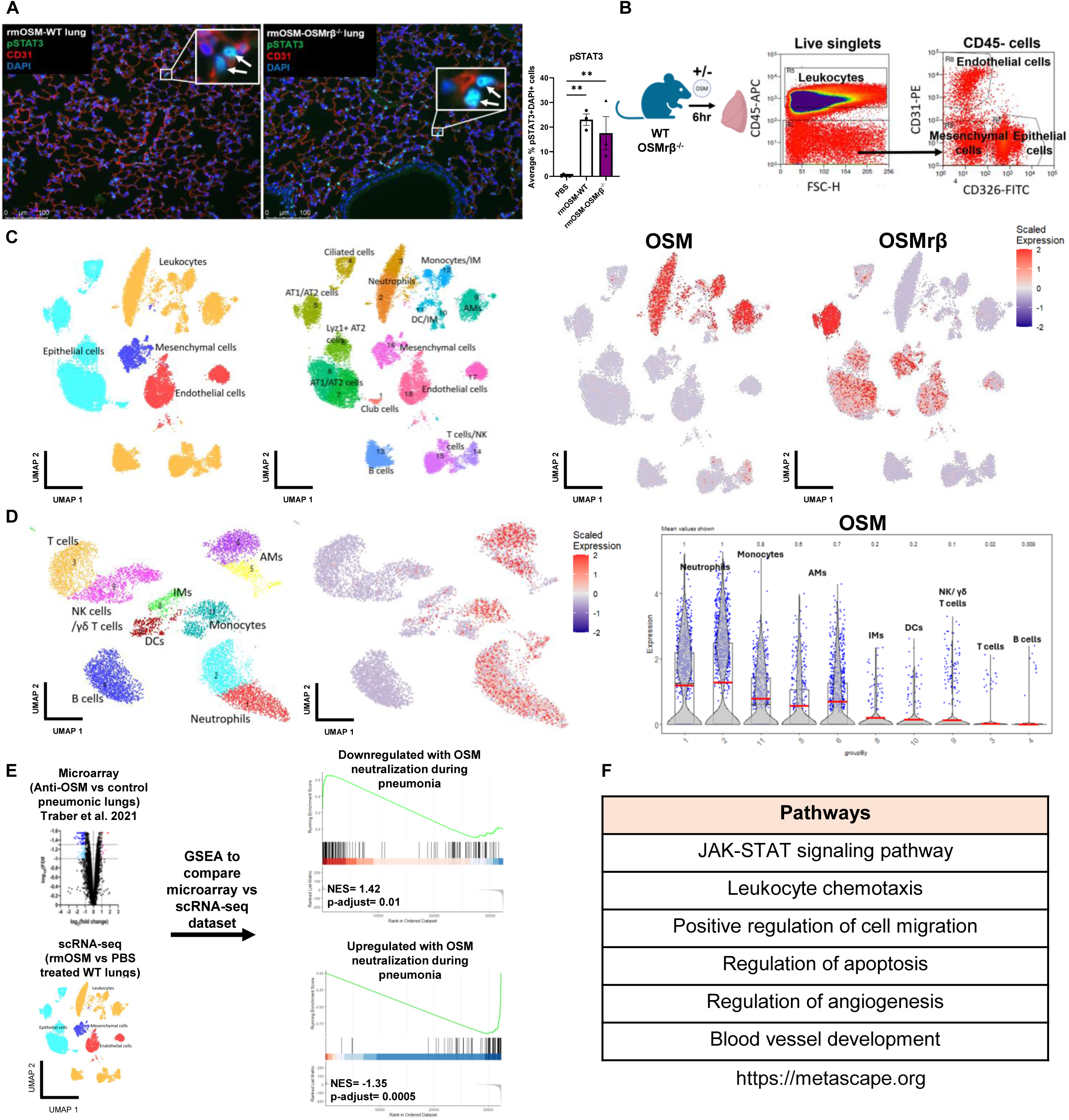
OSM signaling in the lungs occurs both in the presence and absence of OSMrβ. (A) WT and OSMrβ^-/-^ mice were treated intratracheally with rmOSM or vehicle control (PBS). After 1 hour, lungs were collected, fixed, and paraffin embedded. Sections were stained for pSTAT3 (green), CD31 (red), and DAPI (blue) to assess STAT3 activation, and pSTAT3-positive cells were quantified. One-way ANOVA ** P ≤ 0.01. (B) Schematic of the scRNA-sequencing experiment. WT and OSMrβ^-/-^ mice were treated intratracheally with rmOSM or vehicle control (PBS), after which lung leukocyte, epithelial, endothelial, and mesenchymal cell populations were isolated by flow cytometric sorting and processed for scRNA-sequencing. (C) Uniform Manifold Approximation and Projection (UMAP) of annotated cell clusters, with OSM and OSMrβ expression shown across the identified clusters (gene expression is shown on a color scale from low (navy) to high (red)). (D) UMAP of reclustered leukocytes and violin plot of OSM expression across different clusters. (E) Gene set enrichment analysis (GSEA) was conducted to compare previously published microarray data (22) with the scRNA-sequencing dataset. (F) Pathway analysis of OSM-driven genes (FC > 0.1 and FDR < 0.05) identified across microarray and scRNA-sequencing dataset was conducted using Metascape (metascape.org).

To investigate OSMrβ-dependent and independent OSM signaling, we performed transcriptomic analysis of WT and OSMrβ^-/-^ left lobes intratracheally instilled with rmOSM or PBS control for 6 hours. We used flow cytometry to sort leukocytes (CD45^+^), epithelial cells (CD326^+^), endothelial cells (CD31^+^) and mesenchymal cells (CD45^-^CD326^-^CD31^-^), recombined the cell types in equal proportions and performed single-cell RNA sequencing (scRNA-sequencing) on these cells (Figure 4B). After quality control, we utilized standard markers to identify lung cell types (annotations in Table 8 in the Supplementary Tables). Evaluating these cell types, we see OSM exclusively produced by leukocytes and OSMrβ by structural cells, including epithelial cells, endothelial cells and mesenchymal cells (Figure 4C). To look further into what type of leukocytes expressed OSM, we reclustered leukocytes and looked at OSM expression across different types of leukocytes. OSM is highly expressed by certain myeloid cells including neutrophils, monocytes and alveolar macrophages but lowly expressed by interstitial macrophages, dendritic cells and lymphocytes (Figure 4D).

We next compared our scRNA-sequencing to a previously published microarray analysis of *E. coli* infected lungs with and without OSM neutralization.^22^ Gene Set Enrichment Analysis (GSEA) showed that genes that were significantly downregulated with OSM neutralization were upregulated with OSM stimulation (NES=1.42, p-adjusted = 0.01). Likewise, genes that were significantly upregulated with OSM neutralization were downregulated with OSM stimulation (NES=-1.35, p-adjusted = 0.0005) (Figure 4E). Pathway analysis using the web-based platform Metascape^54^ of significantly changed genes in both datasets showed that OSM plays an important role in activating JAK-STAT signaling pathway, regulating leukocyte recruitment and regulating resolution by mediating apoptosis and angiogenesis (Figure 4F).

Using Ingenuity Pathway Analysis (IPA; QIAGEN Inc.), we performed a comparative analysis of OSM signaling across different cell types. We observed that OSMrβ-dependent OSM signaling occurs in epithelial, mesenchymal, and endothelial cells, whereas OSMrβ-independent signaling is restricted to mesenchymal and endothelial cells (Figure 5A). We next examined various cell-type-specific responses to OSM in the presence and absence of OSMrβ. We began by reclustering all epithelial cells and observed that rmOSM treated WT alveolar type 2 (AT2) cells clustered separately from the rest of the AT2 cells (PBS treated WT and OSMrβ^-/-^ and rmOSM treated OSMrβ^-/-^), suggesting that OSMrβ is required for OSM signaling in AT2 cells (Figure 5B). Pathway enrichment analysis by IPA of OSM-treated WT AT2 cells showed increases in type 2 inflammatory responses (“Interleukin-4 and Interleukin-13 signaling”), apoptosis and metabolism (cholesterol biosynthesis, metabolism of polyamines and iron uptake and transport), and downregulation of epithelial adherens junction signaling, PI3K/AKT signaling and RHO GTPase cycle (Figure 5C).

**Figure 5:**
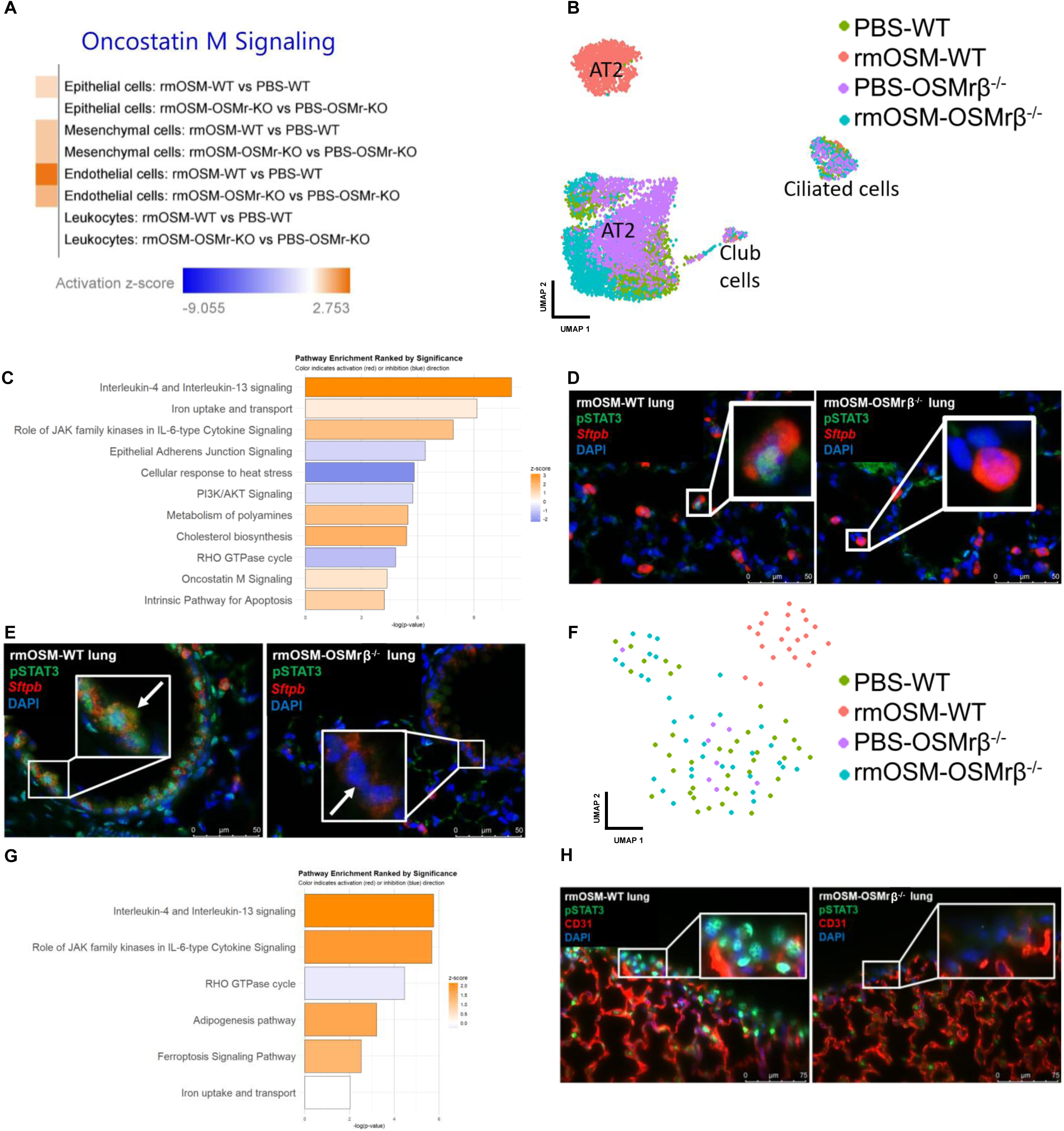
OSMrβ-independent OSM signaling does not occur in lung epithelial cells and mesothelial cells. (A) Pathway analysis using Ingenuity Pathway Analysis (IPA) was performed to compare OSM signaling across differentially expressed gene sets (FDR < 0.05). (B) UMAP representing rmOSM or PBS treated WT or OSMrβ^-/-^ epithelial cells. (C) IPA of rmOSM-treated WT alveolar type II epithelial cells (AT2). Activated pathways are shown in orange, and inhibited pathways are shown in blue. Differentially expressed genes (DEGs) used for IPA were defined by Log₂FC > 0.25 and FDR < 0.05. (D-E) Representative immunofluorescence images of paraffin-embedded lung sections from rmOSM-treated WT and OSMrβ^-/-^ mice, stained for pSTAT3 (green), *Sftpb* (red), and DAPI (blue). AT2 cells were identified as *Sftpb*-positive cells within the alveolar parenchyma, whereas club cells were identified as *Sftpb*-positive cells in the airways. (F) UMAP representing rmOSM or PBS treated WT or OSMrβ^-/-^mesothelial cells. (G) IPA of rmOSM-treated WT mesothelial cells. Activated pathways are shown in orange, and inhibited pathways are shown in blue (Log₂FC > 0.25 and FDR < 0.05). (H) Representative immunofluorescence images of paraffin-embedded lung sections from rmOSM-treated WT and OSMrβ^-/-^ mice, stained for pSTAT3 (green), CD31 (red), and DAPI (blue).

Using immunofluorescence analysis, we confirmed that AT2 cells, which were identified as surfactant protein B (*Sftpb*) positive alveolar cells, respond to OSM by activating STAT3 only in the presence of OSMrβ but not when OSMrβ is absent (Figure 5D). Additionally, club cells, identified as *Sftpb* positive airway cells, respond to OSM by activating STAT3 only in the presence of OSMrβ (Figure 5E).

We next reclustered mesothelial cells and demonstrated that lung mesothelial cells also only respond to OSM in the presence of OSMrβ. We observed that OSM treated WT mesothelial cells clustered together away from the rest of mesothelial cells (Figure 5F). Similar to OSM effect on AT2 cells, OSM activates type 2 inflammatory responses, regulates metabolism, and downregulates RHO GTPase cycle in mesothelial cells (Figure 5G). Using immunofluorescence analysis, we confirmed that OSM activates STAT3 in WT mesothelial cells (identified as cells lining the edge of the lungs) but not OSMrβ^-/-^mesothelial cells (Figure 5H).

Fibroblasts were the most abundant cell type when we reclustered mesenchymal cells. OSM has been shown to be a profibrotic cytokine, especially during chronic disease.^28, 55^ However, we saw a reduction in profibrotic signature with OSM treatment, including decrease in pathways involved in pulmonary fibrosis and collagen production, suggesting that early fibroblast response to OSM involves downregulation of fibrotic pathways (Figure 6A). This finding is consistent with studies suggesting that IL-11, induced downstream of OSM signaling, may be a key driver of fibrosis, rather than OSM itself being directly fibrogenic.^56^ We observed two fibroblast subsets, adventitial fibroblasts (Pdgfra^lo^Ly6a^+^ fibroblasts) and alveolar fibroblasts (Pdgfra^hi^Ly6a^-^ fibroblasts) (Figure 6B). OSM activates WT alveolar fibroblasts and adventitial fibroblasts which were enriched in cluster 1 (Figure 6B). The response of OSMrβ^-/-^ alveolar fibroblasts to OSM differed dramatically from adventitial fibroblasts. OSM stimulation of OSMrβ^-/-^ alveolar fibroblasts resulted in 257 differentially expressed genes (DEGs), comparable to OSM stimulation of WT alveolar fibroblasts, with 388 DEGs. In contrast, OSMrβ^-/-^ adventitial fibroblasts had minimal response to OSM, with 41 DEGs compared to 882 DEGs after OSM stimulation of WT adventitial fibroblasts. Furthermore, OSM-induced chemokine expression was driven predominantly by OSM-treated WT adventitial fibroblasts, whereas alveolar fibroblasts exhibited comparatively lower levels of chemokine expression (Figure 6C). OSM-induced chemokine expression is lost in the absence of OSMrβ in adventitial fibroblasts (Figure 6C).

**Figure 6:**
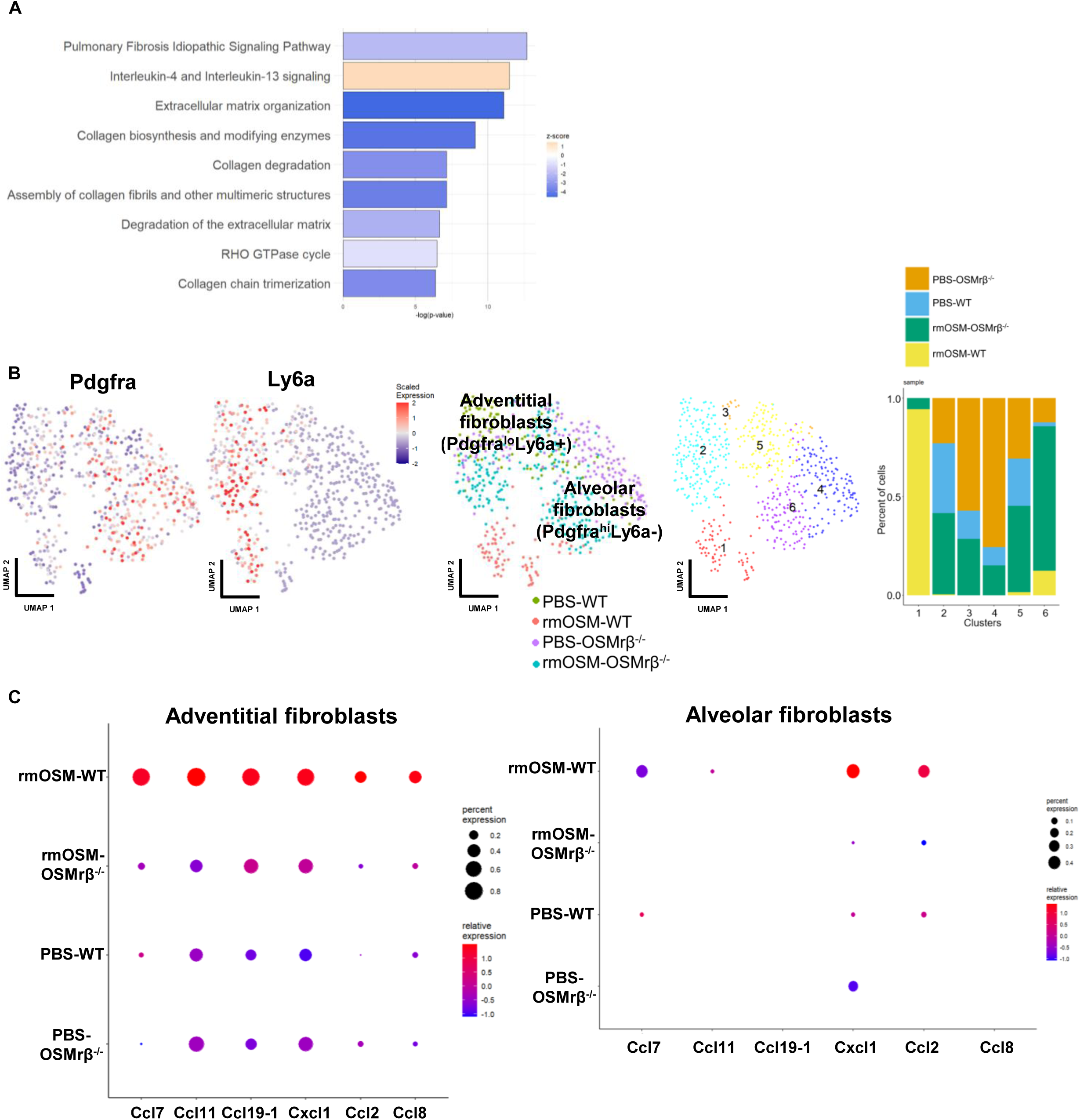
OSM activates both adventitial and alveolar fibroblasts. (A) IPA of rmOSM-treated WT fibroblasts. Activated pathways are shown in orange, and inhibited pathways are shown in blue. DEGs used for IPA were defined by Log₂FC > 0.25 and FDR < 0.05. (B) UMAPs representing Pdgfra and Ly6a expression (gene expression is shown on a color scale from low (navy) to high (red)), UMAP representing rmOSM or PBS treated WT or OSMrβ^-/-^ fibroblasts, UMAP of different fibroblast clusters and stacked bar plot showing the proportional composition of fibroblasts from different samples within each cluster. (C) Dot plot depicting the expression of chemokines in adventitial and alveolar fibroblasts across different samples.

We next extracted fibroblasts from WT and OSMrβ^-/-^ lungs and studied the effect of OSM *ex vivo*. We confirmed that these cells were fibroblasts by ensuring that they were spindle-shaped (typical fibroblast morphology) and that they express fibroblast marker *Col1a1* but not other cell type markers (*Nkx2.1*, *Epcam* and Endothelin-1 (*Edn1*)) (Figure S10A). WT and OSMrβ^-/-^ lung fibroblasts were then treated with rmOSM or vehicle control *ex vivo*. Immunoblot analyses show that OSM activates STAT3 only in the presence of OSMrβ (Figure S10B). Interestingly, OSM seems to regulate MAPK/ERK signaling both in the presence and absence of OSMrβ in fibroblasts, initially decreasing pERK levels (30 minutes post-rmOSM treatment) followed by increase in pERK levels (1 hour post-rmOSM treatment) (Figure S10C).

We next reclustered endothelial cells and observed that amongst lung endothelial cells, general capillary cells (gCaps) from WT and OSMrβ^-/-^ mice treated with rmOSM clustered together away from PBS treated gCaps, suggesting that both OSMrβ-dependent and independent OSM signaling occurs in gCaps (Figure 7A). Pathway analysis showed that OSM activates vascular regeneration in gCaps (mTOR and vascular endothelial growth factor (VEGF) signaling) which is an important function of gCaps,^57^ both in the presence and absence of OSMrβ (Figure 7B). Using RNAscope *in situ* hybridization combined with immunofluorescence analysis, we confirmed that both WT and OSMrβ^-/-^ gCaps, identified as apelin receptor (*Aplnr*) positive alveolar cells, activate STAT3 in response to OSM (Figure 7C). Although we did not capture sufficient arterial and venous endothelial cells to make any meaningful conclusions in our scRNA-sequencing dataset, we observed that OSM activates arterial and venous endothelial cells both in the presence and absence of OSMrβ (Figure 7D).

**Figure 7:**
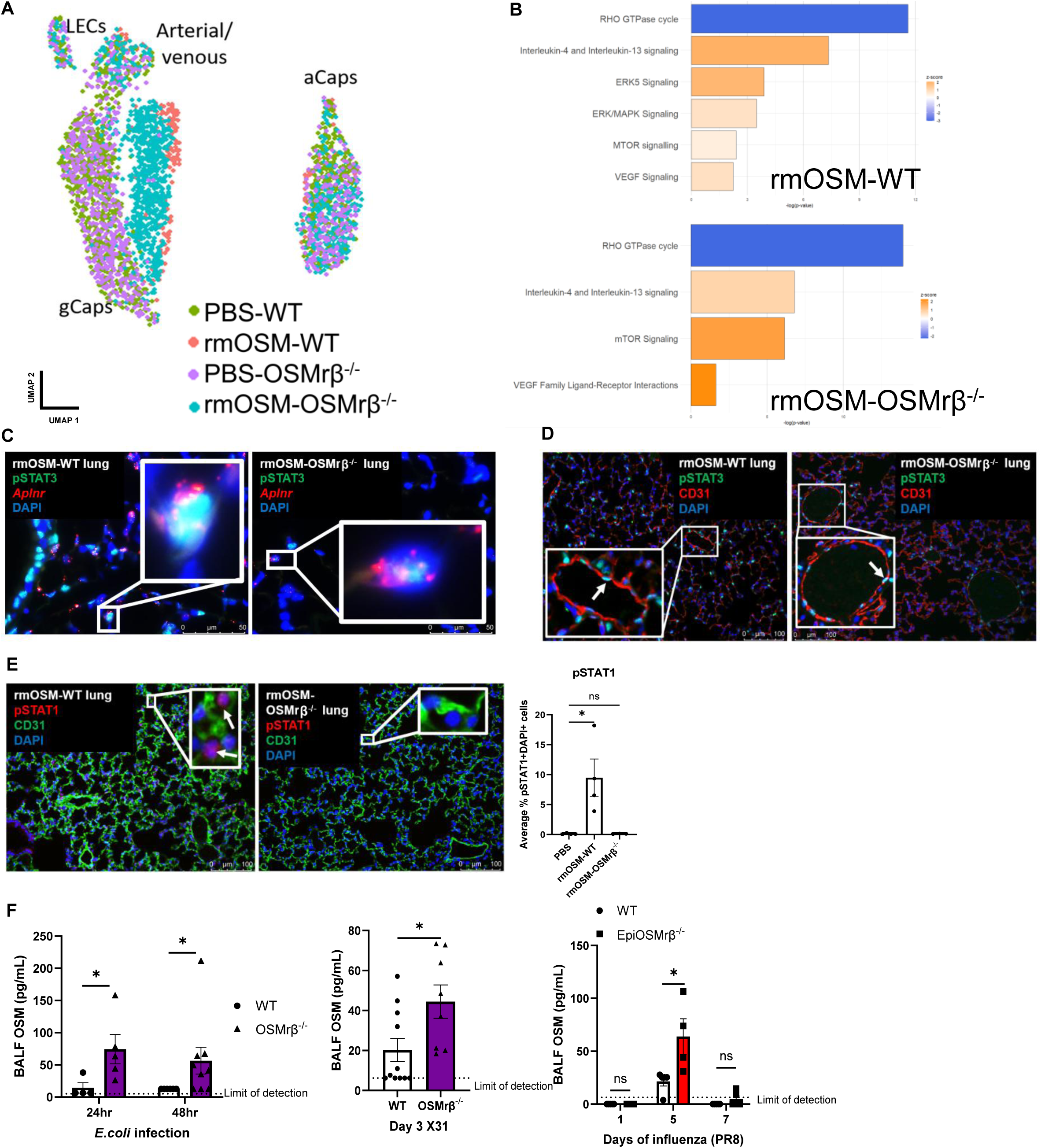
OSMrβ-dependent and independent OSM signaling occur in lung endothelial cells. (A) UMAP representing rmOSM or PBS treated WT or OSMrβ^-/-^ endothelial cells. (B) IPA of rmOSM-treated WT and OSMrβ^-/-^ gCaps. Activated pathways are shown in orange, and inhibited pathways are shown in blue. DEGs used for IPA were defined by Log₂FC > 0.25 and FDR < 0.05. (C) Representative immunofluorescence images of paraffin-embedded lung sections from rmOSM-treated WT and OSMrβ^-/-^ mice, stained for pSTAT3 (green), (C) *Aplnr* (red) or (D) CD31 (red), and DAPI (blue). (E) WT and OSMrβ^-/-^ mice were treated intratracheally with rmOSM or vehicle control (PBS). After 1 hour, lungs were collected, fixed, and paraffin embedded. Sections were stained for pSTAT1 (red), CD31 (green), and DAPI (blue) to assess STAT1 activation, and pSTAT1-positive cells were quantified. One-way ANOVA * P ≤ 0.05. (F) OSM levels in BALF were quantified by ELISA in WT, EpiOSMrβ^-/-^ and OSMrβ^-/-^ lungs infected with *E.coli* or influenza. Mann-Whitney or Multiple Mann-Whitney * P ≤ 0.05. Each data point represents an individual mouse.

The cloning of mouse OSMrβ in 1998 established the prevailing paradigm that mouse OSM signals exclusively via mouse OSMrβ.^29^ More recently however, mouse OSM was shown to signal through mouse LIFrβ to activate STAT3 but not STAT1 in mouse osteoblasts.^32^ To determine whether these findings were recapitulated in the lungs, we also assessed OSM-induced STAT1 activation in the lungs. Consistent with observations in mouse osteoblasts,^32^ OSM induced STAT1 activation in WT lungs but not in OSMrβ^-/-^lungs (Figure 7E). Furthermore, because mouse OSM binds with relatively low affinity to mouse LIFrβ,^32^ higher concentrations of OSM are likely required to elicit biological effects. We measured BALF OSM levels in EpiOSMrβ^-/-^ and OSMrβ^-/-^ mice during influenza and *E. coli* pneumonia and observed increased BALF OSM levels in both genotypes compared with WT controls in both infection models (Figure 7F).

## Discussion

Using mouse models lacking either OSM or OSMrβ, we investigated the importance of OSM-OSMrβ signaling during both bacterial and viral lung infections. Our studies show that OSM is required for protection against both bacterial *E. coli* pneumonia and influenza. Loss of OSM results in increased morbidity, mortality, and cell death. In addition, OSM loss increased inflammatory leukocyte recruitment and a shift of the macrophage population to a more proinflammatory phenotype, which could underlie the increased cell death and mortality seen. Since OSM does not directly act on myeloid cells, we hypothesize that OSM effects on macrophages are indirect.

Remarkably, OSM’s protective effect appears to be OSMrβ-independent, strongly suggesting that OSM signals through an additional, non-canonical receptor. We found that OSM is capable of activating STAT3 signaling depending on the cell type but induces little to no activation of STAT1 in the absence of OSMrβ. Since OSM-LIFrβ signaling has been reported in mouse osteoblasts,^32^ where it similarly drives STAT3 activation without appreciable STAT1 induction, we propose that OSM-LIFrβ signaling is sufficient to rescue the loss of OSM-OSMrβ signaling. LIFrβ is a functional signaling partner of OSM in humans,^28^ and we posit that OSM-mediated protection during pneumonia is largely maintained through LIFrβ-dependent signaling, even in the absence of OSM-OSMrβ signaling. In lung structural cells, LIFrβ is highly expressed by endothelial cells but lowly expressed by club cells and mesothelial cells (Figure S11). We observed that OSM induces STAT3 activation in OSMrβ^-/-^ cells with high expression of LIFrβ (gCaps (Figure 7C) and pulmonary macrovascular endothelial cells (Figure 7D)) but not in OSMrβ^-/-^ cells with low LIFrβ expression (mesothelial cells (Figure 5H) and club cells (Figure 5E)). To our knowledge, this is the first evidence of a critical non-canonical OSM signaling pathway that does not require OSMrβ in the lungs. Investigation of OSM-LIFrβ signaling will be a focus of future investigations.

Recent work has shown that OSM contributes to the restoration of lung epithelial integrity following damage induced by influenza infection and viral mimic-driven inflammatory challenge.^20^ Interestingly, in the work presented here, EpiOSMrβ^-/-^ did not result in any detectable changes in morbidity and lung injury. Furthermore, we found no evidence of OSMrβ-independent OSM signaling in lung epithelial cells, suggesting that direct OSM signaling to epithelial cells is largely dispensable during influenza infection. Therefore, we propose that although OSM can directly stimulate AT2 cell proliferation as previously shown,^20, 44^ its protective effects in influenza can be mediated through alternative receptor signaling pathways, possibly in endothelial cells and alveolar fibroblasts. Nevertheless, a limitation of this study is that our analyses were restricted to peak inflammatory time points and did not assess epithelial regeneration during the recovery phase. Therefore, while epithelial OSMrβ signaling appears dispensable for limiting acute lung injury, its contribution to epithelial regeneration, and restoration of barrier integrity during later stages of infection remains to be determined.

Loss of OSM identified multiple OSM-driven effects on myeloid cells during pneumonia. For example, during *E. coli* pneumonia, loss of OSM does not alter the number of recruited leukocytes, yet we still observed a reduction in AM numbers in the absence of OSM, suggesting that OSM may play an important role in the survival of AMs. In addition, effects of OSM on myeloid cells are mediated through other factors, as myeloid cells do not have OSMrβ.^20, 44^ In the presented data, we have not determined which OSM-induced factors are driving effects on myeloid cells. Potential factors to investigate include those that activate Peroxisome Proliferator-Activated Receptor Gamma (PPAR-γ), a critical transcriptional regulator of AM identity and differentiation.^58, 59, 60^ Given that loss of OSM resulted in delayed monocyte-to-AM differentiation and reduced CD206 expression, both of which are associated with PPAR-γ signaling,^36, 58, 61^ this could be a mediator of OSM effects.

During *E. coli* pneumonia, we observed greater bacteremia and apoptotic lung cells in OSM^-/-^ mice suggesting that OSM is important for protection against injury. GSEA analysis comparing OSM neutralized *E. coli-*infected lungs to OSM stimulated lungs revealed that one of the pathways regulated by OSM is apoptosis. Previous studies have shown that STAT3-mediated anti-apoptotic signaling is important for protection during Respiratory Syncytial Virus (RSV) infection.^62^ Furthermore, IL-6, a cytokine within the same family as OSM, can attenuate RSV-induced epithelial injury through STAT3-dependent anti-apoptotic pathways. Based on these observations, we hypothesize that OSM may similarly contribute to protection against lung injury by engaging STAT3-mediated anti-apoptotic signaling. Consistent with this, OSM potently activates STAT3 but not STAT1 in OSMrβ^-/-^ mice, which are protected from the increased morbidity and cell death observed in OSM^-/-^ mice during pneumonia, suggesting that OSM-induced STAT3 signaling is a key mediator of OSM-mediated protection during lung infection. Future studies will investigate whether OSM-mediated anti-apoptotic pathways contribute to protection against lung injury during bacterial pneumonia and determine which cell types are particularly susceptible to cell death following the loss of OSM signaling.

We observed several discrepancies between the phenotypes of whole-body OSM^-/-^ mice and those previously reported in our OSM neutralization studies during *E. coli* pneumonia.^21, 22^ In our prior work, antibody-mediated neutralization of OSM resulted in increased whole-lung STAT3 and STAT1 activation at 24 hours post-infection.^22^ In contrast, OSM^-/-^ mice showed reduced STAT3 activation and a trend toward reduced STAT1 activation during *E. coli* pneumonia. This discrepancy may be attributable to differences in experimental timing between the two studies. Specifically, STAT activation was assessed at 24 hours post-infection in the neutralization study, whereas analyses in OSM^-/-^ mice were performed at 6 hours post-infection. Given that OSM can induce rapid STAT3 and STAT1 activation within 1 hour of stimulation,^22^ the early response to OSM deficiency is likely reduced STAT signaling. However, by 24 hours post-infection, compensatory mechanisms may be engaged that restore or even augment STAT3 and STAT1 activation, thereby explaining the increased signaling observed in the neutralization model. We will further investigate whether OSM^-/-^ mice similarly exhibit heightened STAT3 and STAT1 activation at later time points during *E. coli* pneumonia.

Previously, we also showed that OSM neutralization reduces BALF neutrophil accumulation during *E. coli* pneumonia.^21^ In the current study, we observed a trend toward reduced extravascular neutrophils in OSM^-/-^ mice compared with WT controls, without reaching statistical significance. This discrepancy is likely due to differences in the experimental approaches used to disrupt OSM signaling. Prior studies have demonstrated that distinct cytokine inhibition strategies can yield divergent phenotypes, in part due to compensatory adaptations that arise in constitutive knockout models.^63, 64^ Whereas antibody-mediated neutralization produces an acute and transient loss of OSM activity, OSM^-/-^ mice lack OSM throughout development and adulthood, which may result in compensatory changes that attenuate acute inflammatory phenotypes. Accordingly, future studies should investigate whether other IL-6 family cytokines, particularly IL-6, LIF, and IL-11, all of which have been implicated in neutrophil recruitment during *E. coli* pneumonia^52, 65, 66^ are upregulated in OSM^-/-^ lungs and contribute to compensatory neutrophil responses.

Another inconsistency between the two models was the degree of bacteremia. While OSM neutralization did not increase bacterial burden in the bloodstream,^21^ OSM^-/-^ mice exhibited elevated bacteremia at 24 hours post-infection. One potential explanation is that the neutralizing antibody may not fully access or inhibit OSM signaling within the vascular compartment and thus preserve endothelial OSM signaling. In contrast, whole-body genetic deletion of OSM would abolish signaling in all compartments, including endothelial cells, potentially impairing vascular barrier integrity and facilitating bacterial dissemination. Supporting this possibility, our scRNA-sequencing data indicates that OSM robustly activates gCaps, a capillary endothelial subset implicated in alveolar capillary repair.^57^ Furthermore, OSM induces pro-angiogenic pathways, including VEGF signaling. Future studies should therefore determine whether endothelial OSM signaling is required for maintaining vascular barrier function and limiting bacterial dissemination during pneumonia.

In conclusion, our work demonstrates that OSM is an important component of the host immune response both during viral and bacterial lung infection. Furthermore, OSMrβ-independent signaling is sufficient to mediate key protective effects of OSM.

## Materials and Methods

### Mice

All genetic knockout mice were bred in an ABSL2 facility. Mice with whole-body deletion of OSMrβ were generated by crossing B6;129-*Osmr^tm1.1Nat^*/J (Strain: 011081, The Jackson Laboratory) with B6.FVB-Tg(EIIa-cre)C5379Lmgd/J (Strain: 003724, The Jackson Laboratory). Mice with epithelial cell-specific deletion of OSMrβ were generated by crossing B6;129-*Osmr^tm1.1Nat^*/J (Strain: 011081, The Jackson Laboratory) with C57BL/6J-Tg(Nkx2-1-cre)2Sand/J (Strain: 008661, The Jackson Laboratory). Mice with whole-body deletion of OSM were generated by first crossing C57BL/6N-Atm1Brd Osmtm1a(KOMP)Wtsi/JMmucd (Stock: 048920-UCD, Mutant Mouse Resource & Research Centers) with B6.129S4-Gt(ROSA)26Sortm1(FLP1)Dym/RainJ (Strain: 009086, The Jackson Laboratory) to delete a portion of the cassette through recombination at FRT sites. Following successful recombination at FRT sites, the homozygous floxed OSM mouse without FLP recombinase were crossed with B6.FVB-Tg(EIIa-cre)C5379Lmgd/J (Strain: 003724, The Jackson Laboratory) to create mice with OSM deletion in all cells. In-house C57BL/6J mice (Strain: 000664, The Jackson Laboratory) that were used as wildtype (WT) controls were co-housed and bred in the same facility as the knockout mice.

### Pneumonia mouse models

Prior to lung infection, 7-13 week-old mice were anesthetized using 75 mg/kg ketamine and 10 mg/kg xylazine via intraperitoneal injection. For influenza infection, mice were infected intratracheally using a 24-gauge angiocatheter with 50μL of vehicle or pathogen into the left lobe of the lungs as previously described. ^67^ Mice were given a sublethal (1000 PFU) or mild (50 PFU) of X31 influenza (influenza A strain/HKx31, H3N2) ^37^ or mild (50 PFU), sublethal (200 PFU) or lethal (400 PFU) dose of PR8 influenza (influenza A strain/Puerto Rico/8/34, H1N1) ^67^ or approximately 1x10^6^ CFU of *E. coli* (serotype 06:K2:H1; ATCC #19138; ATCC).

### Flow cytometry

Mice were anesthetized using 75 mg/kg ketamine and 10 mg/kg xylazine via intraperitoneal injection. To distinguish extravascular leukocytes from intravascular leukocytes, mice were injected retro-orbitally with 100μL of 2μg of anti-CD45.2 flow antibody. The antibodies were allowed to circulate for 3 minutes prior to sacrificing the mice. Lungs were collected in RPMI 1640 media (ThermoFisher Scientific) before processing for flow cytometry. Lungs were digested into single-cell suspensions using digestion buffers containing collagenase type II (Worthington Biochemicals), DNase I (Millipore Sigma) and 125mM calcium chloride. Cells were first incubated with TruStain anti-mouse CD16/32 Fc-block (BioLegend) and True-Stain Monocyte Blocker (BioLegend) prior to staining with flow cytometry antibodies. Flow cytometry was performed on Aurora (Cytek), a high-dimensional multiparameter spectral flow cytometer. SpectraFlo (Cytek) software was used for spectral unmixing of the data and data were analyzed with OMIQ flow cytometry software. The lists of antibodies used in the studies are in Table 1-2.

### Bronchoalveolar lavage collection

Mice were sacrificed using isoflurane overdose and whole lung-heart blocks were removed from the sacrificed mice and tied to a 20-gauge blunted stainless steel catheter via the trachea. Bronchoalveolar lavage fluid (BALF) was collected by instilling and withdrawing ice-cold PBS. BALF was centrifuged at 300g for 5 minutes at 4°C and supernatant was collected for total protein concentration measurement using BCA Protein Assay Kits (ThermoFisher Scientific) and for ELISA to measure proteins of interest. For differential cell counts, the BALF cell pellets were resuspended in ice-cold PBS, counted using LUNA-FL Dual Fluorescence Cell Counter (Logos Biosystems) and diluted to approximately 1x10^6^ cells/mL. 100μL of cell suspension was loaded onto cytocentrifuge funnels and cytocentrifuged onto microscope slides at 800g for 3 minutes (ThermoFisher Scientific Cytospin 4 Centrifuge). Slides were stained with Camco Stain Pak (Fisher Scientific) and percentages of neutrophils, macrophages and lymphocytes were determined to calculate the total number of different types of BALF leukocytes collected from each mouse.

### ELISA

Following the manufacturer’s instruction, DuoSet enzyme-linked immunosorbent assay (R&D Systems) was used to measure mouse OSM in BALF supernatant and whole lung protein homogenates. ELISA was conducted on Nunc MaxiSorp plates (ThermoFisher Scientific) and Synergy LX plate reader (BioTek) was used to read the plates.

### Immunoblotting

Lungs were homogenized using Bullet Blender (Next Advance) according to the manufacturer’s instructions and total protein concentrations were determined using BCA Protein Assay Kits (ThermoFisher Scientific). Immunoblotting was performed as described.^22^ Digital images were quantified using Empiria Studio software (LI-COR) as described.^22^ The list of antibodies used for immunoblotting is in Table 3.

### Histology and immunofluorescence

Following euthanasia of mice using isoflurane overdose, lungs were fixed in 10% Neutral Buffered Formalin (Fisher Scientific) at 23cm H20 pressure. Lungs were fixed for 24-48 hours, dehydrated using increasingly concentrated ethanol, followed by xylene clearing and paraffin embedding. The lung paraffin blocks were cut into 5μm sections for staining. Paraffin-embedded lung sections were deparaffinized in xylene, rehydrated through graded ethanol washes, and subsequently stained with hematoxylin and eosin (H&E), subjected to TUNEL assay, or processed for immunofluorescence staining. Sections stained with H&E staining (Leica) were scored by assessing proportion of the left lung affected by alveolitis, vasculitis, peribroncholitis, proteinaceous edema and airway epithelial denudation.

TUNEL staining was performed using in situ cell death detection kit (Sigma Aldrich) according to the manufacturer’s directions. After TUNEL staining, slides were mounted with VECTASHIELD Vibrance Antifade Mounting Medium with DAPI (Vector Laboratories) and at least 15 images per slide was taken blindly at 200X magnification using a Leica DM4 B light microscope with integrated LAS X software. Quantification of TUNEL-positive cells was done using QuPath software. Briefly, fluorescence images were analyzed in QuPath using the Cell Detection plugin with the DAPI channel used for nuclei detection. TUNEL-positive cells were subsequently identified using an object classifier. The same quantification pipeline was applied to all slides within the experiment using the QuPath Script Editor to ensure consistent analysis across samples.

For immunofluorescence staining, antigen retrieval was performed after deparaffinization of the sections by heating with citrate-based antigen-unmasking solution (Vector Laboratories). Slides were blocked with donkey serum to reduce non-specific antibody binding, and sections were incubated overnight at 4°C with primary antibodies at dilutions mentioned in Table 4. The following day, slides were incubated with secondary antibodies and mounted using VECTASHIELD Vibrance Antifade Mounting Medium with DAPI (Vector Laboratories).

For RNAscope combined with immunofluorescence staining, RNAscope probe hybridization (Advanced Cell Diagnostics) was performed after slide baking and deparaffinization but before immunofluorescence staining according to the manufacturer’s protocol (RNAscope Multiplex Fluorescent Reagent Kit v2). Image acquisition and quantification were performed using a similar approach as described for the TUNEL analysis mentioned above. The list of antibodies used for immunofluorescence is in Table 4 and the list of RNAscope probes used is in Table 5.

### Pathogen burden assessment

Following euthanasia of mice using isoflurane overdose, lungs and blood were collected for pathogen burden assessment. For viral plaque assays, the left lungs from influenza-infected mice were minced into small pieces using a razor blade in a petri dish and subsequently homogenized using a Bullet Blender (Next Advance). The homogenates were centrifuged at 5,000g for 5 minutes at 4°C, and the supernatants were collected. Ten-fold serial dilutions of the supernatants were prepared and used to infect MDCK (Madin–Darby Canine Kidney) cells. Plaques were counted 3 days post-infection. Furthermore, RNA was isolated from a portion of the remaining lung homogenate supernatant, and X31 matrix viral RNA levels were quantified by RT-qPCR.

For measurement of lung bacterial burden, the left lungs from *E. coli*-infected mice were homogenized using a Bullet Blender and serially diluted ten-fold in water. A total of 100 µL of each dilution was plated onto 5% sheep’s blood agar plates (Fisher Scientific) and incubated overnight at 37°C. Colony-forming units (CFUs) were enumerated the following day. For bacteremia measurements, blood was collected from the inferior vena cava (IVC) of the mice using a heparinized syringe and serially diluted ten-fold in water. A total of 100 µL of each dilution was plated onto 5% sheep’s blood agar plates and incubated overnight at 37°C. CFUs were enumerated the following day.

### RNA isolation and RT-qPCR

The left lungs were harvested and homogenized in RLT buffer (Qiagen) supplemented with β-mercaptoethanol using a Bullet Blender (Next Advance). For cultured cells, cell lysates were prepared directly in RLT buffer containing β-mercaptoethanol. Total RNA was isolated according to the manufacturer’s instructions using the RNeasy Mini Kit (Qiagen) for lung tissue or the RNeasy Micro Kit (Qiagen) for cultured cells. RNA concentration and purity were determined using a NanoDrop spectrophotometer (ThermoFisher Scientific).

Gene expression was quantified by RT-qPCR using the TaqMan RNA-to-CT 1-Step Kit and a QuantStudio 3 Real-Time PCR System (ThermoFisher Scientific). Threshold cycle (Ct) values for target genes were normalized to 18S rRNA as the endogenous control.

### Single-cell RNA sequencing

WT and OSMrβ^-/-^ mice were intratracheally instilled with recombinant mouse OSM (rmOSM) or vehicle control (PBS). Six hours later, the left lungs were harvested and enzymatically digested in a solution containing elastase, collagenase II, dispase (Worthington Biochemical), and DNase I (MilliporeSigma). Single-cell suspensions were incubated with TruStain anti-mouse CD16/32 Fc-block (BioLegend) and True-Stain Monocyte Blocker (BioLegend) prior to staining with flow cytometry antibodies. Leukocytes (live CD45^+^ cells), epithelial cells (live CD45^-^ EpCAM^+^ CD31^-^ cells), endothelial cells (live CD45^-^ EpCAM^-^ CD31^+^ cells), and mesenchymal cells (live CD45^-^EpCAM^-^ CD31^-^ cells) were sorted using a FACSAria II SORP cell sorter (BD Biosciences).

Sorted cell populations were combined in equal proportions and processed for single-cell RNA sequencing. The list of antibodies used for flow sorting is in Table 6. The sorted cells were processed using the 10x Genomics Chromium (GemCode) platform, in which individual cells, reagents, and barcoded Gel Beads were encapsulated into nanoliter-scale Gel Bead-in-Emulsions (GEMs). Within each GEM, cells were lysed and RNA underwent barcoded reverse transcription. Full-length barcoded cDNA was subsequently amplified by PCR to generate sufficient material for library preparation. Single-cell RNA-sequencing libraries were generated by the Boston University Department of Medicine Single Cell Sequencing Core Facility and library size distribution and concentration were assessed using a Bioanalyzer High Sensitivity DNA Assay (Agilent Technologies). Libraries were sequenced on an Illumina NextSeq 2000 according to the manufacturer’s protocols (Illumina and 10x Genomics).

Comprehensive quality control (QC) of the single-cell RNA-sequencing data was performed using Cell Ranger package (10x Genomics) and SCTK-QC pipeline.^68^ The single-cell RNA-sequencing QC metrics are summarized in Table 7. Cells with fewer than 500 UMIs, fewer than 500 detected genes, or greater than 5% mitochondrial transcripts were excluded. The number of cells for each sample are 4,918, 6,319, 6,831 and 6,635 for TK_LY_M1 (rmOSM treated WT lung), TK_LY_M2 (PBS treated WT lung), TK_LY_M3 (rmOSM treated OSMrβ^-/-^ lung) and TK_LY_M4 (PBS treated OSMrβ^-/-^ lung) respectively. A total of 24,703 cells were used for downstream analyses. The single-cell RNA sequencing data will be deposited in the NCBI Gene Expression Omnibus (GEO).

Sequencing data was analyzed and visualized in R (version 4.3.1) using Rstudio (R packages and versions include celda 1.18.2, SingleCellExperiment 1.24.0, ggplot2 3.5.2, dittoSeq 1.14.3). Differentially expressed genes (DEGs) with a log₂fold change (log₂FC) > 0.25 and a false discovery rate (FDR) < 0.05 were used for Ingenuity Pathway Analysis (IPA; QIAGEN) and Metascape pathway enrichment analysis. Cell populations in the single-cell RNA sequencing dataset were annotated using established marker genes and previously published cell-type annotations, ^50, 69, 70^ as described in Table 8.

### Lung fibroblast extraction and studies

Primary mouse lung fibroblasts were isolated from freshly harvested WT and OSMrβ^-/-^lungs by mechanical dissociation and enzymatic digestion with collagenase II and dispase (Worthington Biochemical) in DMEM/F12 media. Digested tissue was washed, centrifuged at 524g for 5 minutes at 4°C, and resuspended in complete DMEM/F12 media before plating in 10-cm culture dishes. Fibroblasts were allowed to migrate from tissue explants and adhere to culture plates over 7 days at 37°C in a humidified incubator with 5% CO₂, with media changes and transfer of tissue fragments to fresh plates as needed. After 7 days, tissue fragments were removed and fibroblasts were replated in EMEM media (15% FBS, penicillin/streptomycin, non-essential amino acids, and sodium pyruvate) to enrich for fibroblast growth. Fibroblast identity was confirmed by RT-qPCR analysis of RNA isolated from cultured cells, using established fibroblast marker genes.

### Statistical analysis

Statistical analyses were performed using GraphPad Prism (version 10). Data normality was determined using a Shapiro-Wilk test. Normally distributed datAassa were analyzed using parametric tests (t-test or ANOVA). Non-normally distributed data were analyzed using non-parametric tests (Mann-Whitney or Kruskal-Wallis) or log-transformed followed by analysis using an appropriate parametric test. Data were presented as means ± SEM. Comparisons were considered significant at P ≤ 0.05.

## Acknowledgements

Funding sources:

KET: R01HL158732, KL2TR001411, K08KL130582, UL1TR001439

LJQ: R01HL165718, R21AI191650

MPV: F32HL178206

JPM: R01HL171499, R01AI162850 MRJ: R01HL164612

MB: 1R01HL166588, 1R01HL181357, R01HL181444

We would like to thank the Flow Cytometry Core (Anna C. Belkina, Brian R. Tilton and Shari C. Brezinsky) and the Single Cell Sequencing Core (Salam H. AI Abdullatif and Yuriy Alekseyev) of Boston University Chobanian & Avedisian School of Medicine for their expert assistance.

## References

1. Traber, K.E. & Mizgerd, J.P. The Integrated Pulmonary Immune Response to Pneumonia. Annu Rev Immunol 43, 545–569 (2025).

2. Torres, A. et al. Pneumonia. Nat Rev Dis Primers 7, 25 (2021).

3. Cilloniz, C., Dela Cruz, C.S., Dy-Agra, G., Pagcatipunan, R.S., Jr. & Pneumo-Strategy, G. World Pneumonia Day 2024: Fighting Pneumonia and Antimicrobial Resistance. Am J Respir Crit Care Med 210, 1283–1285 (2024).

4. Dela Cruz, C.S., et al. Understanding the Host in the Management of Pneumonia. An Official American Thoracic Society Workshop Report. Ann Am Thorac Soc 18, 1087–1097 (2021).

5. Aleem, M.S., Sexton, R. & Akella, J. Pneumonia in an Immunocompromised Patient. StatPearls: Treasure Island (FL), 2026.

6. Smith, D.K., Kuckel, D.P. & Recidoro, A.M. Community-Acquired Pneumonia in Children: Rapid Evidence Review. Am Fam Physician 104, 618–625 (2021).

7. Jain, S. et al. Community-Acquired Pneumonia Requiring Hospitalization among U.S. Adults. N Engl J Med 373, 415–427 (2015).

8. Khan, M.A., Bajwa, A. & Hussain, S.T. Pneumonia: Recent Updates on Diagnosis and Treatment. Microorganisms 13 (2025).

9. Zhang, H. et al. Antiviral treatment for viral pneumonia: current drugs and natural compounds. Virol J 22, 62 (2025).

10. Dela Cruz, C.S., et al. Future Research Directions in Pneumonia. NHLBI Working Group Report. Am J Respir Crit Care Med 198, 256–263 (2018).

11. Quinton, L.J., Walkey, A.J. & Mizgerd, J.P. Integrative Physiology of Pneumonia. Physiol Rev 98, 1417–1464 (2018).

12. Wolf, C.L., Pruett, C., Lighter, D. & Jorcyk, C.L. The clinical relevance of OSM in inflammatory diseases: a comprehensive review. Front Immunol 14, 1239732 (2023).

13. Ayaub, E.A. et al. Overexpression of OSM and IL-6 impacts the polarization of pro-fibrotic macrophages and the development of bleomycin-induced lung fibrosis. Sci Rep 7, 13281 (2017).

14. Wong, S., Botelho, F.M., Rodrigues, R.M. & Richards, C.D. Oncostatin M overexpression induces matrix deposition, STAT3 activation, and SMAD1 Dysregulation in lungs of fibrosis-resistant BALB/c mice. Lab Invest 94, 1003-1016 (2014).

15. Simpson, J.L., Baines, K.J., Boyle, M.J., Scott, R.J. & Gibson, P.G. Oncostatin M (OSM) is increased in asthma with incompletely reversible airflow obstruction. Exp Lung Res 35, 781–794 (2009).

16. Headland, S.E. et al. Oncostatin M expression induced by bacterial triggers drives airway inflammatory and mucus secretion in severe asthma. Sci Transl Med 14, eabf8188 (2022).

17. Shien, K. et al. JAK1/STAT3 Activation through a Proinflammatory Cytokine Pathway Leads to Resistance to Molecularly Targeted Therapy in Non-Small Cell Lung Cancer. Mol Cancer Ther 16, 2234–2245 (2017).

18. Lauber, S. et al. Novel function of Oncostatin M as a potent tumour-promoting agent in lung. Int J Cancer 136, 831–843 (2015).

19. Cichy, J. & Pure, E. Oncostatin M and transforming growth factor-beta 1 induce post-translational modification and hyaluronan binding to CD44 in lung-derived epithelial tumor cells. J Biol Chem 275, 18061–18069 (2000).

20. Hoagland, D.A. et al. Macrophage-derived oncostatin M repairs the lung epithelial barrier during inflammatory damage. Science 389, 169–175 (2025).

21. Traber, K.E. et al. Induction of STAT3-Dependent CXCL5 Expression and Neutrophil Recruitment by Oncostatin-M during Pneumonia. Am J Respir Cell Mol Biol 53, 479–488 (2015).

22. Traber, K.E. et al. Neutrophil-Derived Oncostatin M Triggers Diverse Signaling Pathways during Pneumonia. Infect Immun 89 (2021).

23. Quinton, L.J. et al. Alveolar epithelial STAT3, IL-6 family cytokines, and host defense during Escherichia coli pneumonia. Am J Respir Cell Mol Biol 38, 699-706 (2008).

24. West, N.R., Owens, B.M.J. & Hegazy, A.N. The oncostatin M-stromal cell axis in health and disease. Scand J Immunol 88, e12694 (2018).

25. Dawson, R.E., Jenkins, B.J. & Saad, M.I. IL-6 family cytokines in respiratory health and disease. Cytokine 143, 155520 (2021).

26. Heinrich, P.C. et al. Principles of interleukin (IL)-6-type cytokine signalling and its regulation. Biochem J 374, 1–20 (2003).

27. MacDonald, K., Botelho, F., Ashkar, A.A. & Richards, C.D. Type I Interferon Signaling is Required for Oncostatin-M Driven Inflammatory Responses in Mouse Lung. J Interferon Cytokine Res 42, 568–579 (2022).

28. Stawski, L. & Trojanowska, M. Oncostatin M and its role in fibrosis. Connect Tissue Res 60, 40–49 (2019).

29. Lindberg, R.A. et al. Cloning and characterization of a specific receptor for mouse oncostatin M. Mol Cell Biol 18, 3357–3367 (1998).

30. Adrian-Segarra, J.M. et al. The AB loop of oncostatin M (OSM) determines species-specific signaling in humans and mice. J Biol Chem 293, 20181-20199 (2018).

31. Walker, E.C. et al. Oncostatin M promotes bone formation independently of resorption when signaling through leukemia inhibitory factor receptor in mice. J Clin Invest 120, 582–592 (2010).

32. Walker, E.C. et al. Murine Oncostatin M Acts via Leukemia Inhibitory Factor Receptor to Phosphorylate Signal Transducer and Activator of Transcription 3 (STAT3) but Not STAT1, an Effect That Protects Bone Mass. J Biol Chem 291, 21703–21716 (2016).

33. Askovich, P.S. et al. Differential host response, rather than early viral replication efficiency, correlates with pathogenicity caused by influenza viruses. PLoS One 8, e74863 (2013).

34. Iwasaki, A. & Pillai, P.S. Innate immunity to influenza virus infection. Nat Rev Immunol 14, 315–328 (2014).

35. Short, K.R., Kroeze, E., Fouchier, R.A.M. & Kuiken, T. Pathogenesis of influenza-induced acute respiratory distress syndrome. Lancet Infect Dis 14, 57–69 (2014).

36. David, C., Verney, C., Si-Tahar, M. & Guillon, A. The deadly dance of alveolar macrophages and influenza virus. Eur Respir Rev 33 (2024).

37. Aegerter, H. et al. Influenza-induced monocyte-derived alveolar macrophages confer prolonged antibacterial protection. Nat Immunol 21, 145–157 (2020).

38. Li, F., et al. Monocyte-derived alveolar macrophages autonomously determine severe outcome of respiratory viral infection. Sci Immunol 7, eabj5761 (2022).

39. Ghoneim, H.E., Thomas, P.G. & McCullers, J.A. Depletion of Alveolar Macrophages during Influenza Infection Facilitates Bacterial Superinfections. J Immunol 191, 1250–1259 (2013).

40. Ruscitti, C., Radermecker, C. & Marichal, T. Journey of monocytes and macrophages upon influenza A virus infection. Curr Opin Virol 66, 101409 (2024).

41. Hou, F., Xiao, K., Tang, L. & Xie, L. Diversity of Macrophages in Lung Homeostasis and Diseases. Front Immunol 12, 753940 (2021).

42. Han, H. et al. Oncostatin M promotes infarct repair and improves cardiac function after myocardial infarction. Am J Transl Res 13, 11329–11340 (2021).

43. Sims, N.A. & Levesque, J.P. Oncostatin M: Dual Regulator of the Skeletal and Hematopoietic Systems. Curr Osteoporos Rep 22, 80–95 (2024).

44. Shang, X. et al. Monocyte-derived macrophages support alveolar regeneration via oncostatin M post-H1N1 infection during the recovery phase. Respir Res 26, 285 (2025).

45. Dey, G. et al. Signaling network of Oncostatin M pathway. J Cell Commun Signal 7, 103–108 (2013).

46. Schyns, J. et al. Non-classical tissue monocytes and two functionally distinct populations of interstitial macrophages populate the mouse lung. Nat Commun 10, 3964 (2019).

47. Vanneste, D. et al. MafB-restricted local monocyte proliferation precedes lung interstitial macrophage differentiation. Nature immunology 24, 827–840 (2023).

48. Gu, Y., Lawrence, T., Mohamed, R., Liang, Y. & Yahaya, B.H. The emerging roles of interstitial macrophages in pulmonary fibrosis: A perspective from scRNA-seq analyses. Front Immunol 13, 923235 (2022).

49. Duggan, J.M. et al. Synergistic interactions of TLR2/6 and TLR9 induce a high level of resistance to lung infection in mice. J Immunol 186, 5916–5926 (2011).

50. Soucy, A.M., et al. Transcriptomic responses of lung mesenchymal cells during pneumonia. JCI Insight 10 (2025).

51. Broug-Holub, E. et al. Alveolar macrophages are required for protective pulmonary defenses in murine Klebsiella pneumonia: elimination of alveolar macrophages increases neutrophil recruitment but decreases bacterial clearance and survival. Infect Immun 65, 1139–1146 (1997).

52. Quinton, L.J. et al. Leukemia Inhibitory Factor Signaling Is Required for Lung Protection during Pneumonia. J Immunol 188, 6300–6308 (2012).

53. Na, E. et al. Epithelial LIF signaling limits apoptosis and lung injury during bacterial pneumonia. Am J Physiol Lung Cell Mol Physiol 322, L550–L563 (2022).

54. Zhou, Y. et al. Metascape provides a biologist-oriented resource for the analysis of systems-level datasets. Nat Commun 10, 1523 (2019).

55. Mozaffarian, A. et al. Mechanisms of oncostatin M-induced pulmonary inflammation and fibrosis. J Immunol 181, 7243–7253 (2008).

56. Ng, B. et al. Interleukin-11 is a therapeutic target in idiopathic pulmonary fibrosis. Sci Transl Med 11 (2019).

57. Gillich, A. et al. Capillary cell-type specialization in the alveolus. Nature 586, 785–789 (2020).

58. Smith, M.R., Standiford, T.J. & Reddy, R.C. PPARs in alveolar macrophage biology. PPAR Res 2007, 23812 (2007).

59. Baker, A.D. et al. Targeted PPARgamma deficiency in alveolar macrophages disrupts surfactant catabolism. J Lipid Res 51, 1325–1331 (2010).

60. Huang, S. et al. PPAR-gamma in Macrophages Limits Pulmonary Inflammation and Promotes Host Recovery following Respiratory Viral Infection. J Virol 93 (2019).

61. Schneider, C. et al. Induction of the nuclear receptor PPAR-gamma by the cytokine GM-CSF is critical for the differentiation of fetal monocytes into alveolar macrophages. Nat Immunol 15, 1026–1037 (2014).

62. Zhao, C. et al. Activation of STAT3-mediated ciliated cell survival protects against severe infection by respiratory syncytial virus. J Clin Invest 134 (2024).

63. El-Brolosy, M.A. & Stainier, D.Y.R. Genetic compensation: A phenomenon in search of mechanisms. PLoS Genet 13, e1006780 (2017).

64. Velasco-Aviles, S. et al. A genetic compensatory mechanism regulated by Jun and Mef2d modulates the expression of distinct class IIa Hdacs to ensure peripheral nerve myelination and repair. Elife 11 (2022).

65. Jones, M.R. et al. Roles of interleukin-6 in activation of STAT proteins and recruitment of neutrophils during Escherichia coli pneumonia. J Infect Dis 193, 360–369 (2006).

66. Traber, K.E. et al. Roles of interleukin-11 during acute bacterial pneumonia. PloS one 14, e0221029 (2019).

67. Crossey, E. et al. Influenza induces lung lymphangiogenesis independent of YAP/TAZ activity in lymphatic endothelial cells. Sci Rep 14, 21324 (2024).

68. Hong, R. et al. Comprehensive generation, visualization, and reporting of quality control metrics for single-cell RNA sequencing data. Nat Commun 13, 1688 (2022).

69. Aegerter, H., Lambrecht, B.N. & Jakubzick, C.V. Biology of lung macrophages in health and disease. Immunity 55, 1564–1580 (2022).

70. Hurskainen, M. et al. Single cell transcriptomic analysis of murine lung development on hyperoxia-induced damage. Nat Commun 12, 1565 (2021).

