## Supplementary Figures for "Non-canonical Oncostatin M Signaling Provides Protection during Respiratory Viral and Bacterial infections"

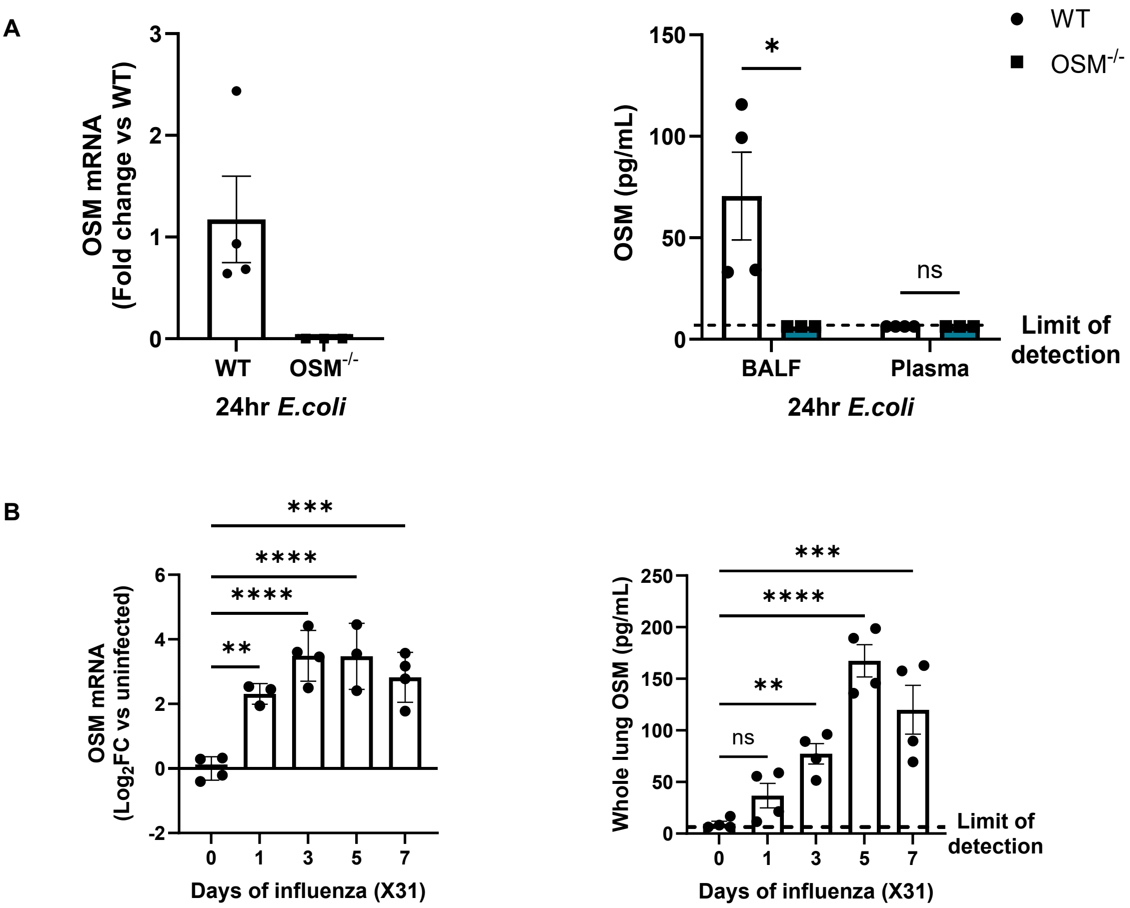


**Figure S1: Validation of OSM$^{-/-}$ model and OSM measurement during influenza.** (A) WT and OSM$^{-/-}$ mice were intratracheally infected with *E.coli* and lung tissue was collected at 24hr post-infection for quantification of OSM RNA using RT-qPCR. BALF from the lungs was collected to measure OSM protein level using ELISA. (B) WT mice were intratracheally infected with 1000 PFU of X31 (influenza A/HKx31, H3N2) and lung tissue was collected at baseline (day 0) and at days 1, 3, 5, and 7 post-infection for quantification of OSM RNA (RT-qPCR) and protein (ELISA) expression. Two-way ANOVA for (A). One-way ANOVA for (B) * $P \leq 0.05$, ** $P \leq 0.01$, *** $P \leq 0.001$ and **** $P < 0.0001$. Each data point represents an individual mouse.

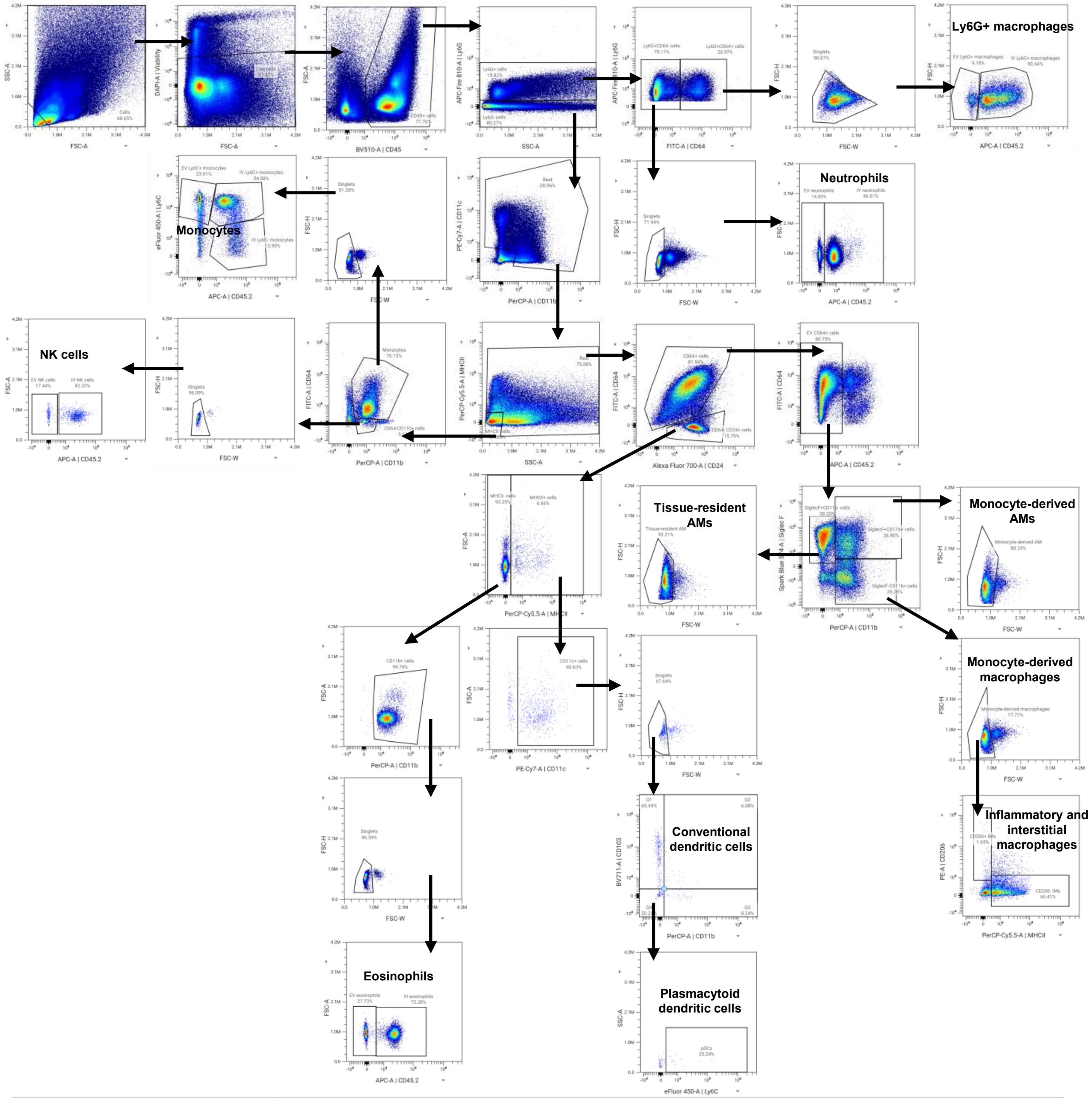


| | |
|---|---|
| **Neutrophils** | **Live CD45+ Ly6G+ CD64-** |
| **Ly6G+ macrophages** | **Live CD45+ Ly6G+ CD64+** |
| **Tissue-resident AMs (TRAMs)** | **Live CD45+ Ly6G- EVCD64+ SiglecF+ CD11b-** |
| **Monocyte-derived AMs (moAMs)** | **Live CD45+ Ly6G- EVCD64+ SiglecF+ CD11b+** |
| **Inflammatory and interstitial macrophages** | **Live CD45+ Ly6G- EVCD64+ SiglecF- CD11b+ with either CD206+ MCHII- or CD206- MHCII+** |
| **Monocytes** | **Live CD45+ Ly6G- MHCII- CD64+ CD11b+ Ly6C+** |
| **NK cells** | **Live CD45+ Ly6G- MHCII- CD64- CD11b+** |
| **Eosinophils** | **Live CD45+ Ly6G- CD64- CD24+ MHCII- CD11b+** |
| **Conventional dendritic cells** | **Live CD45+ Ly6G- CD64- CD24+ MHCII+ CD11c+ CD103+/- CD11b+/-** |
| **Plasmacytoid dendritic cells** | **Live CD45+ Ly6G- CD64- CD24+ MHCII+ CD11c+ CD103- CD11b- Ly6C+** |

**Figure S2: Gating strategies for flow cytometry analysis of lung leukocytes during X31 influenza infection.** Flow cytometry gating strategy adapted from Supplementary Figure 2 of Yu et al (DOI: 10.1371/journal.pone.0150606).

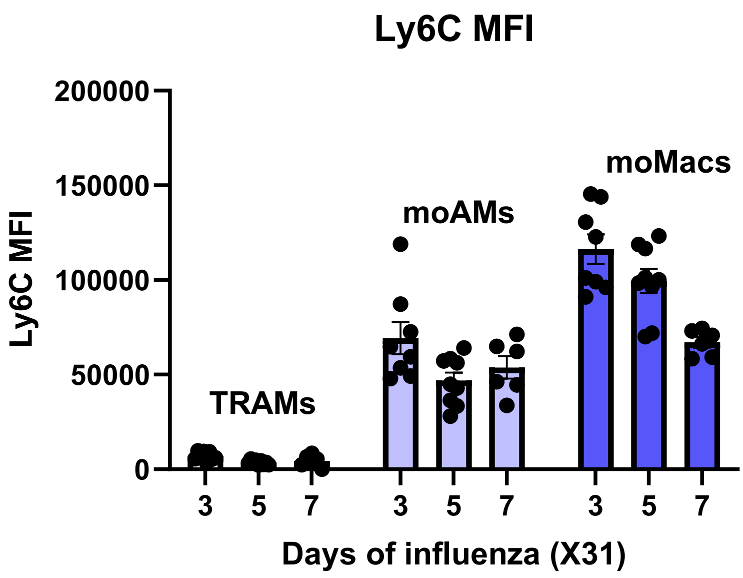


**Figure S3: Ly6C Median Fluorescence Intensity expression of myeloid cells.** Ly6C expression on WT tissue-resident AMs (TRAMs), monocyte-derived AMs (moAMs) and monocyte-derived macrophages (moMacs) at day 3, day 5 and day 7 post-influenza infection.

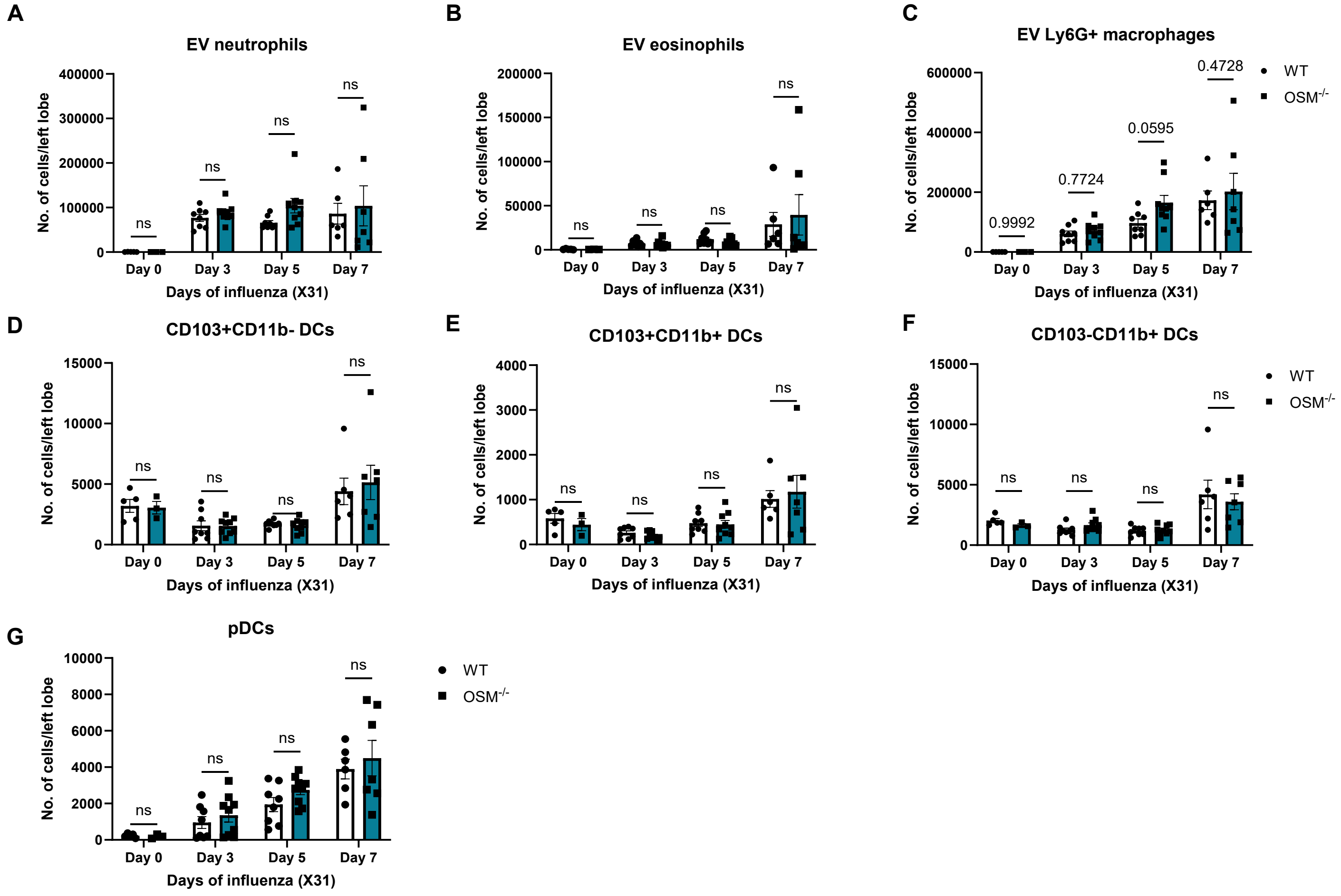


**Figure S4: Flow cytometric quantification of indicated leukocyte populations during X31 influenza infection.** Leukocytes in the left lung of WT and OSM$^{-/-}$ mice harvested at baseline (day 0), day 3, day 5 and day 7 post X31 infection were quantified by spectral flow cytometry. Mice were intravascularly instilled with CD45.2 flow antibody to distinguish extravascular leukocytes from intravascular leukocytes. Absolute cell counts of EV (IV CD45.2-) (A) neutrophils (B) eosinophils (C) Ly6G+ macrophages (D-F) conventional dendritic cells (CD103+CD11b-, CD103+CD11b+, CD103-CD11b+) and (G) plasmacytoid dendritic cells. At least two independent experiments. Two-way ANOVA. Each data point represents an individual mouse.

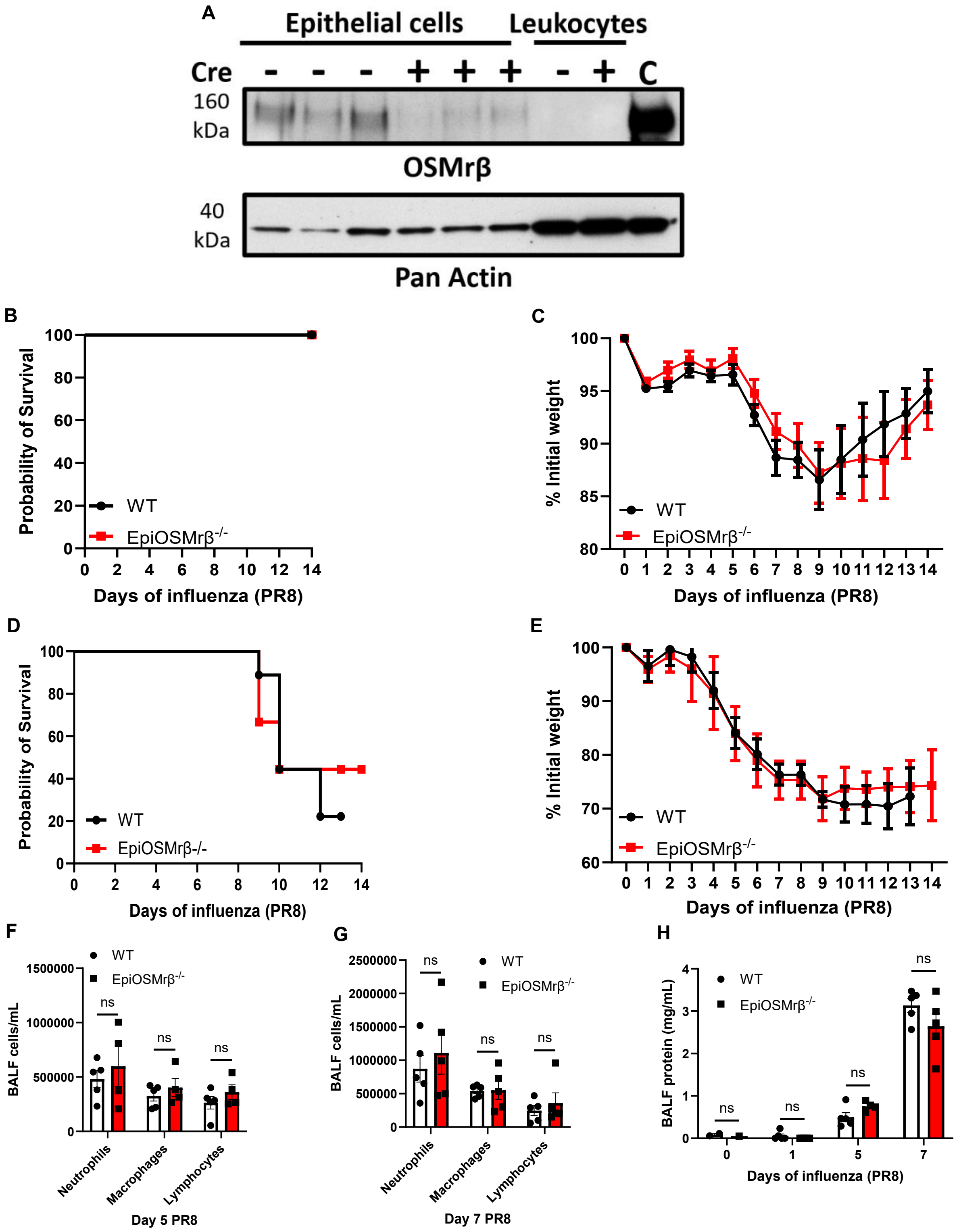


**Figure S5: OSMrβ on lung epithelial cells is dispensable for OSM-mediated protection during influenza.** (A) OSMrβ protein measured from epithelial cells sorted from Cre+ and Cre- lungs from EpiOSMrβ$^{-/-}$ mice. Leukocytes sorted from Cre+ and Cre- lungs from EpiOSMrβ$^{-/-}$ mice and whole lung protein C were used as control. (B) Survival curve and (C) weights of WT and EpiOSMrβ$^{-/-}$ mice that were intratracheally infected with sublethal dose of 50 PFU of PR8 influenza (n=9 for WT and n=10 for EpiOSMrβ$^{-/-}$). (D) Survival curve and (E) weights of WT and EpiOSMrβ$^{-/-}$ mice that were intratracheally infected with lethal dose of 400 PFU of PR8 influenza (n=9 for WT and n=9 for EpiOSMrβ$^{-/-}$). Log-rank (Mantel-Cox) test for (B) and (D). Two-way ANOVA for (C) and (E). Number of neutrophils, macrophages and lymphocytes in bronchoalveolar lavage fluid (BALF) in lungs at (F) 5 days and (G) 7 days post-infection. (H) Total protein in BALF from lungs at baseline (day 0), day 1, day 5 and day 7 post-infection. Two-way ANOVA. Each data point represents an individual mouse.

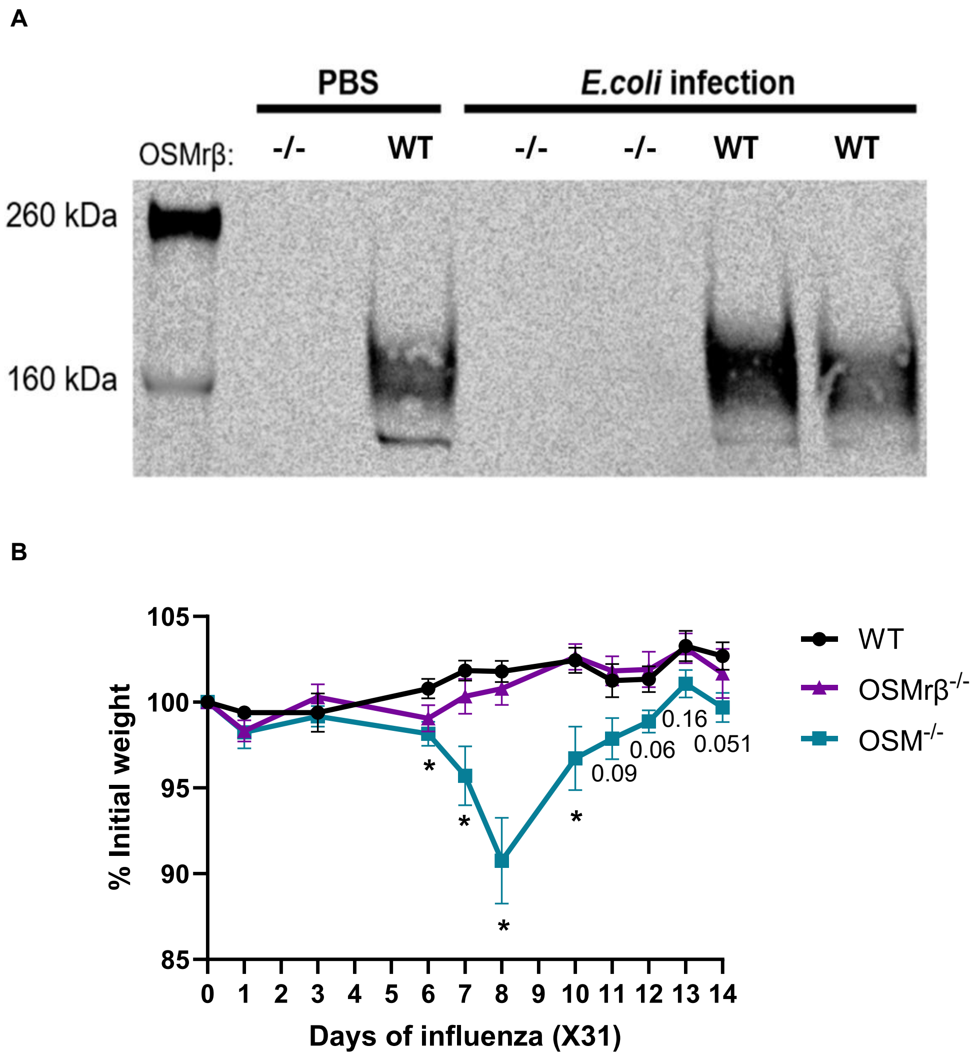


**Figure S6: OSMrβ-independent OSM signaling is sufficient to mediate protection during influenza infection.** (A) Protein was extracted from WT and OSMrβ$^{-/-}$ lungs 24 hours after intratracheal instillation of either PBS or *E. coli*. Immunoblot analysis was then performed to assess OSMrβ protein levels. (B) Weights of WT, OSMrβ$^{-/-}$ and OSM$^{-/-}$ mice that were intratracheally infected with 50 PFU of X31 influenza (n=6 for WT, n=9 for OSMrβ$^{-/-}$ and n=6 for OSM$^{-/-}$). Two-way ANOVA * $P \leq 0.05$.

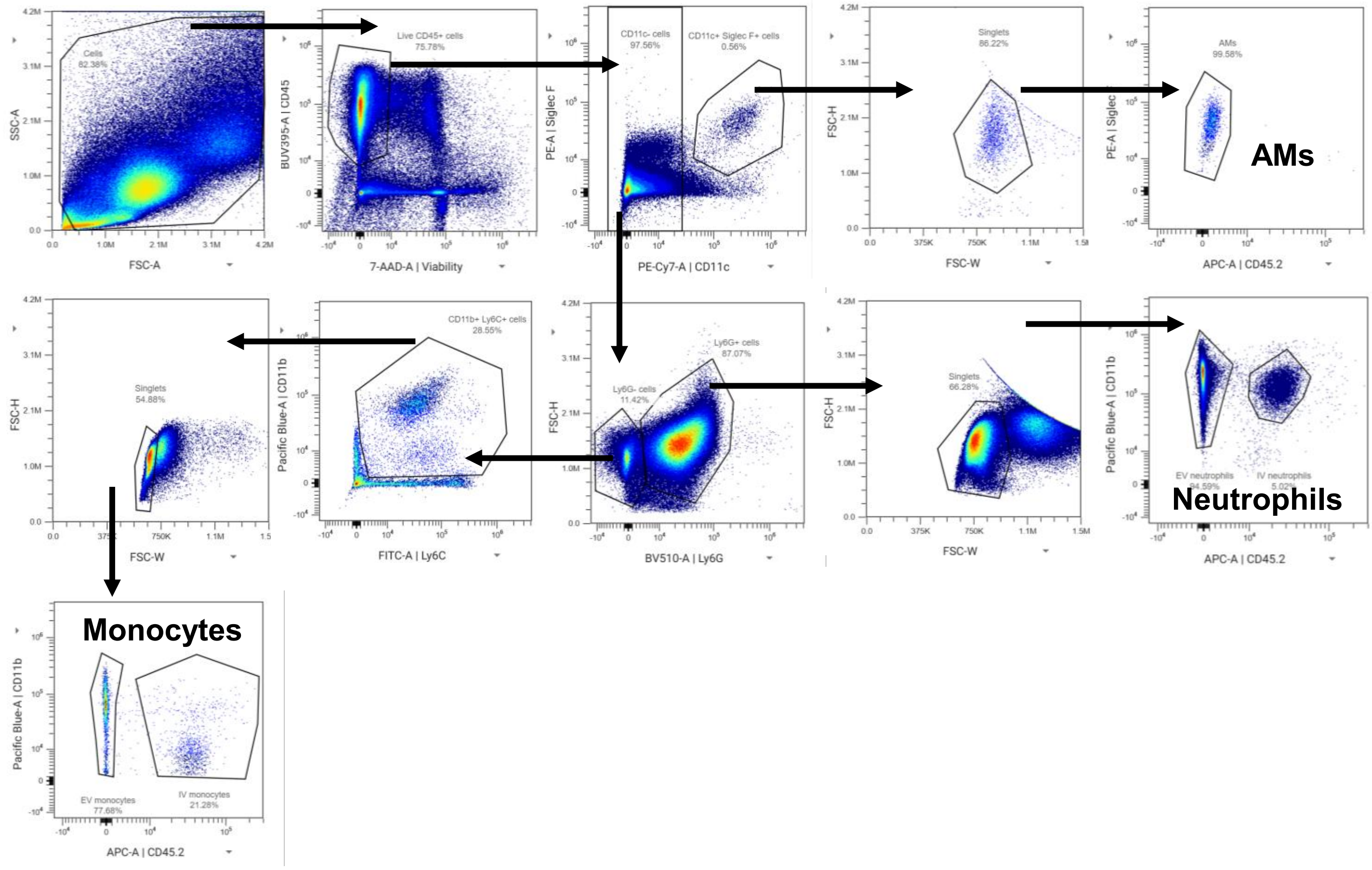


| | |
|---|---|
| **Neutrophils** | Live CD45+ CD11c- Ly6G+ |
| **Monocytes** | Live CD45+ CD11c- Ly6G- CD11b+ Ly6C+ |
| **Alveolar macrophages (AMs)** | CD45+ CD11c+ SiglecF+ |

**Figure S7: Gating strategies for flow cytometry analysis of lung leukocytes during *E.coli* pneumonia.**

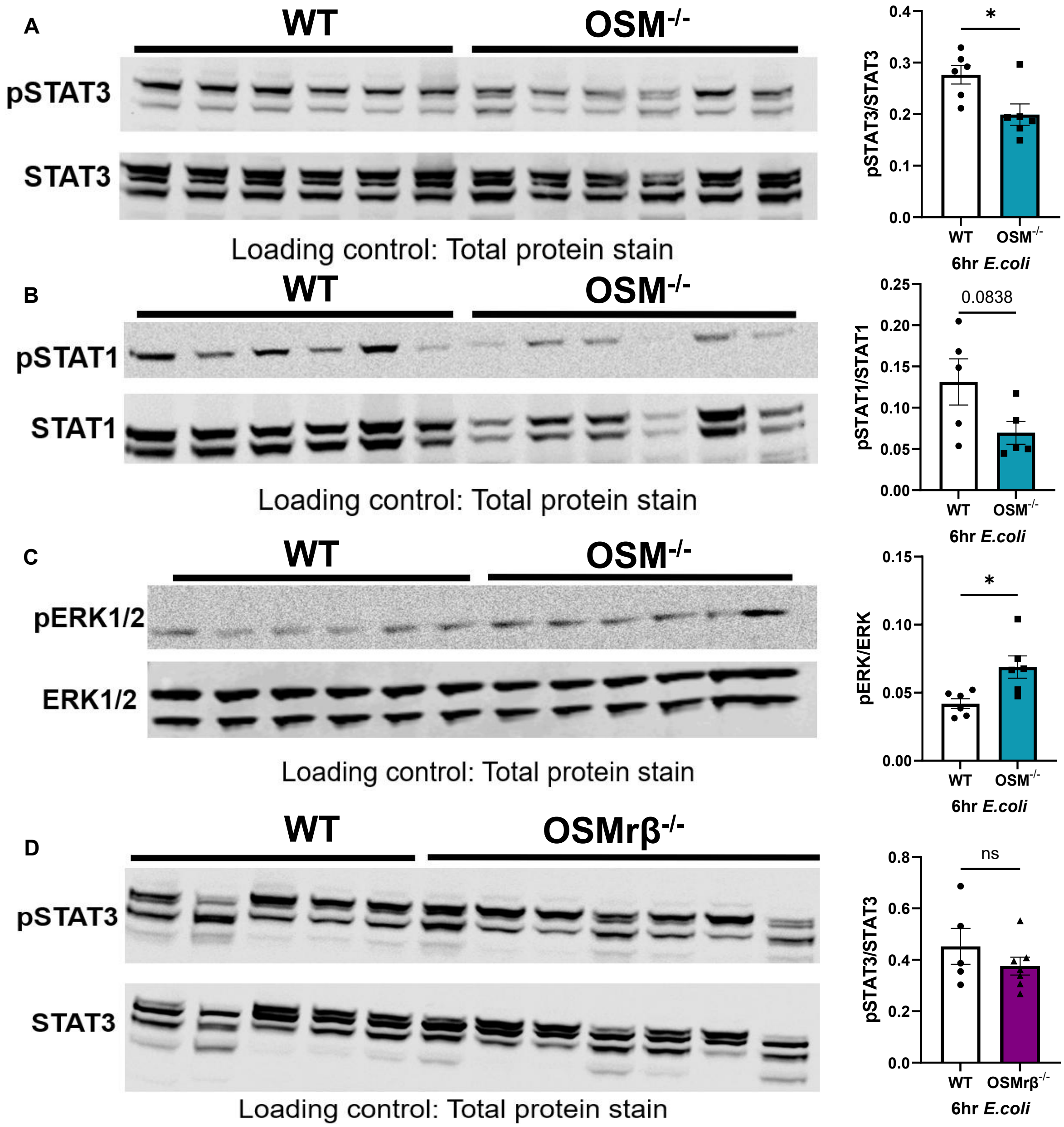


**Figure S8: Loss of OSM but not OSMrβ leads to changes in lung transcription activation during *E.coli* pneumonia.** WT, $OSM^{-/-}$ and $OSMr\beta^{-/-}$ mice were intratracheally infected with *E.coli* and left lungs were collected 6hrs post-infection. Immunoblot analysis of (A) pSTAT3 and total STAT3 (B) pSTAT1 and total STAT1 (C) pERK and total ERK of WT and $OSM^{-/-}$ lung protein. (D) pSTAT3 and total STAT3 of WT and $OSMr\beta^{-/-}$ lung protein. Total protein staining was used as a loading control. Female mice were used for experiments in this figure. Unpaired T-test * $P \leq 0.05$. Each data point represents an individual mouse.

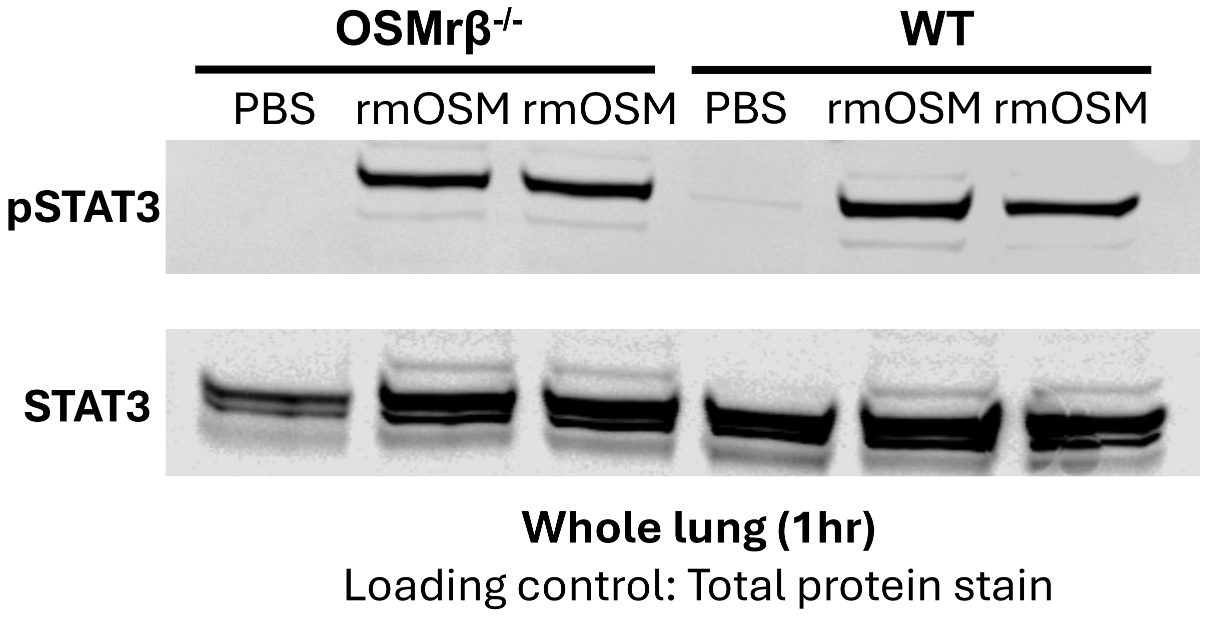


**Figure S9: OSM activates STAT3 in the lungs both in the presence and absence of OSMrβ.**
WT and OSMrβ$^{-/-}$ mice were intratracheally instilled with rmOSM or PBS and left lungs were collected 1 hour later. Immunoblot analysis of pSTAT3 and total STAT3 from lung protein. Total protein staining was used as a loading control.

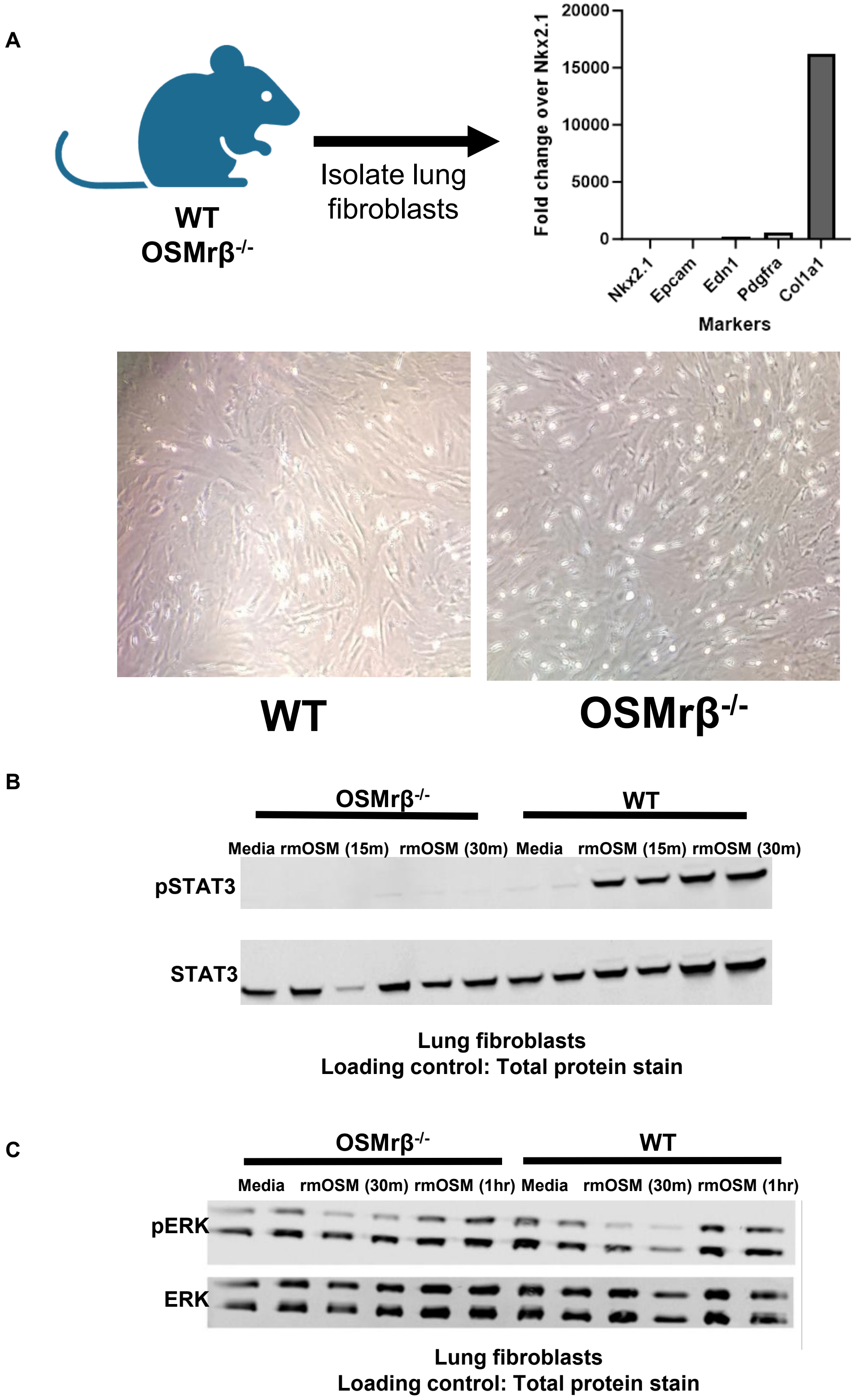


**Figure S10: Isolation and characterization of primary lung fibroblasts**. (A) Fibroblasts were isolated from WT and OSMrβ$^{-/-}$ lungs by mechanical and enzymatic digestion and enriched through culture in EMEM medium. Expressions of Nkx2.1, Epcam, Edn1, Pdgfra, and Col1a1 were measured by RT-qPCR using RNA extracted from cultured fibroblasts. Representative brightfield images of fibroblasts in culture are shown. Immunoblot analysis of (B) pSTAT3 and total STAT3, and (C) pERK and total ERK in fibroblasts isolated from WT or OSMrβ$^{-/-}$ lungs following *ex vivo* treatment with rmOSM or media. Total protein staining was used as a loading control.

### LIFrβ

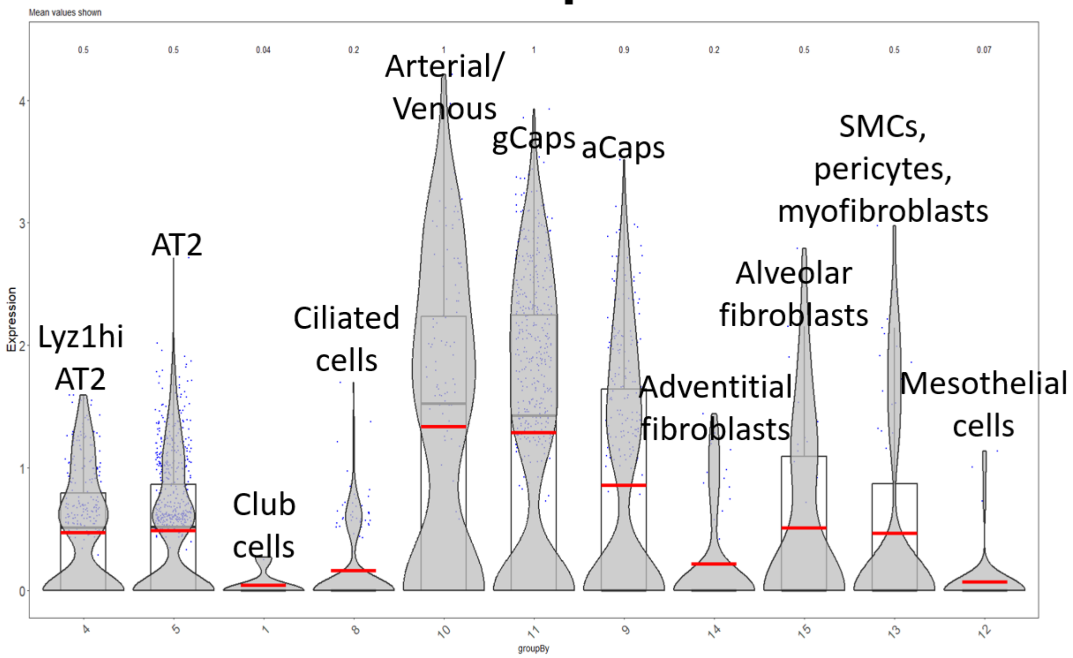


**Figure S11: LIFrβ expression.** Violin plot showing LIFrβ expression across different lung structural cell populations.
