## Supplementary Tables for "Non-canonical Oncostatin M Signaling Provides Protection during Respiratory Viral and Bacterial infections"

**Table 1: Myeloid flow panel for X31 influenza infected lungs.**

| **Antibodies** | **Source** | **Catalog #** | **Dilution** |
|---|---|---|---|
| BV510 CD45 (Clone: 30-F11) | BD Biosciences | 563891 | 1:400 |
| APC CD45.2 (Clone: 104) | BioLegend | 109814 | 1:10 |
| PerCP CD11b (Clone: M1/70) | BioLegend | 101230 | 1:100 |
| PE-Cy7 CD11c (Clone: HL3) | BD Biosciences | 558079 | 1:1600 |
| Alexa Fluor 700 CD24 (Clone: M1/69) | BioLegend | 101835 | 1:200 |
| FITC CD64 (Clone: X54-5/7.1) | BioLegend | 139316 | 1:200 |
| eFluor 450 Ly6C (Clone: HK1.4) | Invitrogen | 48-5932-82 | 1:800 |
| PerCP-Cy5.5 MHCII (Clone: M5/114.15.2) | BD Biosciences | 562363 | 1:800 |
| BV711 CD103 (Clone: 2E7) | BioLegend | 121435 | 1:100 |
| PE CD206 (Clone: C068C2) | BioLegend | 141706 | 1:800 |
| Spark Blue 574 CD170 (SiglecF) (Clone: S17007L) | BioLegend | 285184 | 1:200 |
| APC-Fire810 Ly6G (Clone: 1A8) | BioLegend | 127669 | 1:3200 |
| BV785 F4/80 (Clone: BM8) | BioLegend | 123141 | 1:200 |
| BUV395 CD274 (PDL1) (Clone: MIH5) | BD Biosciences | 745616 | 1:100 |
| **Viability** | **Source** | **Catalog #** | **Dilution** |
| NucBlue Live Cell Stain ReadyProbes reagent | Invitrogen | R37605 | 1:2000 |

**Table 2: Myeloid flow panel for *E. coli* infected lungs.**

| **Antibodies** | **Source** | **Catalog #** | **Dilution** |
|---|---|---|---|
| BUV395 CD45 (Clone: 30-F11) | BD Biosciences | 564279 | 1:500 |
| APC CD45.2 (Clone: 104) | BioLegend | 109814 | 1:10 |
| Pacific Blue CD11b (Clone: M1/70) | BioLegend | 101223 | 1:600 |
| PE-Cy7 CD11c (Clone: N418) | BioLegend | 117317 | 1:800 |
| BV510 Ly6G (Clone: 1A8) | BioLegend | 127633 | 1:500 |
| FITC Ly6C (Clone: HK1.4) | BioLegend | 128005 | 1:600 |
| PE SiglecF (Clone: E50-2440) | BD Biosciences | 552126 | 1:800 |
| **Viability** | **Source** | **Catalog #** | **Dilution** |
| 7-AAD Viability Staining | BioLegend | 420404 | 1:100 |

**Table 3: Immunoblotting antibodies.**

| Antibodies | Source | Catalog # | Dilution |
|---|---|---|---|
| Phospho-Stat3 (Tyr705) (D3A7) Rabbit mAb | Cell signaling | 9145 | 1:1000 |
| STAT3 (D3Z2G) Rabbit mAb | Cell signaling | 12640 | 1:1000 |
| Phospho-Stat1 (Ser727) Rabbit pAb | Cell signaling | 9177 | 1:1000 |
| Stat1 Rabbit pAb | Cell signaling | 9172 | 1:1000 |
| Phospho-p44/42 MAPK (Erk1/2) (Thr202/Tyr204) (D13.14.4E) Rabbit mAb | Cell signaling | 4370 | 1:1000 |
| p44/42 MAPK (Erk1/2) (137F5) Rabbit mAb | Cell signaling | 4695 | 1:1000 |
| OSM receptor Goat pAb | R&D systems | AF662 | 1:1000 |
| Pan Actin (D18C11) Rabbit mAb | Cell signaling | 8456 | 1:1000 |

**Table 4: Immunofluorescence antibodies.**

| Antibodies | Source | Catalog # | Dilution |
|---|---|---|---|
| Phospho-Stat3 (Tyr705) (D3A7) Rabbit mAb | Cell signaling | 9145 | 1:100 |
| Phospho-Stat1 (Ser727) Rabbit pAb | Cell signaling | 9177 | 1:100 |
| CD31/PECAM-1 Goat pAb | R&D systems | AF3628 | 3:100 |
| CSF-1R/M-CSF-R (E6W9F) Rabbit mAb | Cell signaling | 43390 | 1:100 |

**Table 5: RNAscope probes.**

| Probes | Source | Catalog # |
|---|---|---|
| Probe-Mm-Aplnr | ACDBio | 436171 |
| Probe-Mm-Sftpb-No-XHs-C2 | ACDBio | 539421-C2 |

**Table 6: Flow panel for rmOSM or PBS treated lungs for flow sorting.**

| Antibodies | Source | Catalog # | Dilution |
|---|---|---|---|
| APC CD45 (Clone: I3/2.3) | BioLegend | 147707 | 1:500 |
| PE CD31 (Clone: 390) | BioLegend | 102407 | 1:100 |
| FITC CD326 (Epcam) (Clone: G8.8) | BioLegend | 118207 | 1:200 |
| **Viability** | **Source** | **Catalog #** | **Dilution** |
| 7-AAD Viability Staining | BioLegend | 420404 | 1:100 |
| Calcein blue AM | ThermoFisher Scientific | C1429 | 1:200 |

**Table 7: Single-cell RNA-sequencing quality control (QC) metrics.**

| Sample | Number of reads | Mean reads per cell | Median UMI counts per cell | Median genes per cell | Total genes detected | Median % of mitochondrial counts | Median ambient RNA DecontX contamination score |
|---|---|---|---|---|---|---|---|
| TK_LY_M1 (rmOSM treated WT lung) | 219,230,034 | 44,577 | 4,857 | 1,790 | 21,026 | 4.12% | 0.0097 |
| TK_LY_M2 (PBS treated WT lung) | 289,609,541 | 45,832 | 6,387 | 2,353 | 21,967 | 5.01% | 0.0079 |
| TK_LY_M3 (rmOSM treated OSMrβ$^{-/-}$ lung) | 301,090,990 | 44,077 | 5,034 | 1,858 | 22,134 | 4.08% | 0.0079 |
| TK_LY_M4 (PBS treated OSMrβ$^{-/-}$ lung) | 348,102,098 | 52,465 | 5,765 | 1,944 | 22,002 | 3.72% | 0.0075 |

**Table 8: Cell type annotations for single-cell RNA sequencing data.**

| Cell type | Marker genes |
|---|---|
| *All leukocytes identified as Ptprc⁺ cells* | |
| Neutrophils | Ly6g, Itgam, Ly6c2, G0s2, Asprv1 |
| Alveolar macrophages (AMs) | Chil3, Atp6v0d2, Lpl, Ctsd, Marco, Siglecf, Itgax, Fcgr1 |
| B cells | Igkc, Cd74, Ly6d, Cd79a, Ms4a1, Cd19 |
| NK cells | S100a10, Lgals1, Ccl5, Aw112010, Nkg7, |
| γδ T cells | S100a4, Cxcr6, Trdc, S100a10, Lgals1, Ikzf2 |
| T cells | Ly6c2, Nkg7, Cd8a, Klrd1, Cd8b1, Lef1, Ms4a6b, Tcf7 |
| Monocytes | Plac8, Ifitm3, Ifi2712a, Pou2f2, S100a4, Apoe, Ccr2, Vcan, C5ar1 |
| Interstitial macrophages (IMs) | Pf4, C1qa, C1qb, C1qc, Apoe, Mafb, C5ar1 |
| Dendritic cells (DCs) | Itgae, Epsti1, Traf1, Fscn1, Cd74, H2-Eb1, H2-Aa, H2-Ab1, Cst3 |
| *All endothelial cells identified as Pecam1⁺ cells* | |
| Lymphatic endothelial cells (LECs) | Mmrn1, Ccl21a-1, Fgl2, Nrp2, Nts, Pdpn, Reln |
| Arterial endothelial cells | Vwf, Slc6a2, Car8, Eln, Fabp4, Fbln2 |
| Venous endothelial cells | Mgp, Fbln5, Ltbp4, Eln, Vwf, Fbln2, Cdh13 |
| General capillary cells (gCaps) | Kit, Gpihbp1, Itga6, Peg3, Cd93 |
| Aerocytes (aCaps) | Emp2, Car4, Igfbp7, Ednrb |
| *All epithelial cells identified as Epcam⁺ cells* | |
| Alveolar type II cells (AT2s) | Lyz2, Sftpc, Slc34a2, S100g, Lamp3, Abca3, Sftpa1 |
| Club cells | Scgb1a1, Scgb3a2, Reg3g, Scgb3a1 |
| Ciliated cells | Foxj1, Tmem212, Dynlrb2, Cdkn1c, Tppp3 |
| *Mesothelial cells identified as Msln⁺ cells* | |
| Mesothelial cells | Msln, Igfbp5, Igfbp6, Upk3b, Rarres2, Crip1 |
| *All mesenchymal cells identified as Ptprc-Pecam1-Epcam- cells* | |
| Alveolar fibroblasts | Mfap4, Macf1, Npnt, Limch1, Gyg, Co113a1, Pdgfra |
| Adventitial fibroblasts | Dcn, Mfap5, Col1a1, Col1a2, Col14a1, Ly6a |
